# Lasting effects of low-calorie sweeteners on glucose regulation, sugar intake, and memory

**DOI:** 10.1101/2021.11.22.469487

**Authors:** Linda Tsan, Sandrine Chometton, Yanning Zuo, Shan Sun, Anna M. R. Hayes, Lana Bridi, Rae Lan, Anthony A. Fodor, Emily E. Noble, Xia Yang, Scott E. Kanoski, Lindsey A. Schier

**Affiliations:** Neuroscience Graduate Program, University of Southern California, Los Angeles, CA, USA; Department of Biological Sciences, Human and Evolutionary Biology Section, University of Southern California, Los Angeles, CA, USA; Department of Integrative Biology and Physiology, University of California at Los Angeles, Los Angeles, CA, USA; Department of Bioinformatics and Genomics at the University of North Carolina at Charlotte, Charlotte, NC, USA; Department of Nutritional Sciences, University of Georgia, Athens, GA, USA

## Abstract

Low-calorie sweetener (LCS) consumption in children has increased due to widespread LCS presence in the food environment and efforts to mitigate obesity through sugar replacement. However, mechanistic studies on the impact of early-life LCS consumption are lacking. Therefore, we developed a rodent model to evaluate the effects of daily LCS consumption (acesulfame potassium, saccharin, or stevia) during adolescence on adult metabolic, gut microbiome, neural, and behavioral outcomes. Results reveal that habitual early-life LCS consumption disrupts post-oral glucose tolerance and impairs hippocampal-dependent memory in the absence of weight gain. Furthermore, LCS consumption reduces lingual sweet taste receptor expression and alters sugar-motivated appetitive and consummatory responses. RNA sequencing analyses reveal that LCS also impacts collagen- and synaptic signaling-related gene pathways in the hippocampus and nucleus accumbens, respectively, in a sex-dependent manner. Collectively, these results suggest that regular early-life LCS consumption yields long-lasting impairments in metabolism, sugar-motivated behavior, and hippocampal-dependent memory.

## 1. Introduction

Children are the highest sugar consumers of any age group with approximately 16% of total calories coming from added sugar, nearly 40% of which comes from sugar-sweetened beverages (SSBs) (*1*). Studies in humans and rodent models reveal that excessive early life sugar consumption negatively impacts glucose metabolism (*2*, *3*), sensory/reward processing (*4*, *5*), the gut microbiome (*6*, *7*), and neurocognitive abilities (*8*–*11*). To combat these adverse effects, children, like adults, are consuming more foods and beverages that contain low calorie sweeteners (LCS) in lieu of sugar (*12*, *13*). Indeed, LCS consumption in children increased by nearly 200% between 1999 and 2012 (*12*). Even though substitution of sugar-laden foods and fluids with those containing LCS is perceived to be beneficial for body weight management, the evidence for this in youth is mixed (*14*–*20*). Mechanistic rodent model studies may offer some insight into the long-term effects of habitual early life LCS consumption on caloric intake and metabolic function.

Although LCS bind to the same taste receptors as caloric sugars (*21*), they are less effective at eliciting biological signals that influence food intake and metabolism. For example, carbohydrates and other nutrients trigger the release of hormones from intestinal enteroendocrine cells that influence satiation and glucose metabolism, including glucagon-like-peptide 1 (GLP-1) and glucose-dependent insulinotropic polypeptide (GIP) (*22*). LCS consumption, however, fails to stimulate GLP-1 secretion in rats (*23*) and can lead to over-secretion of GIP in humans (*24*). Some have suggested that extended LCS consumption results in the uncoupling of sweet taste from the physiological and neural events normally produced by sugars, which, over time, degrades the ability of sweet taste cues to effectively guide food choice, satiety, and metabolic processes (*25*). Consistent with this view, a history of LCS consumption in rodent models attenuates cephalic phase physiological signaling (*26*, *27*).

In addition to physiological metabolic outcomes, emerging evidence indicates that LCS consumption may be linked with neurocognitive function. For example, habitual LCS consumption based on food-frequency questionnaires is associated with increased prospective risk for all-cause dementia^28^. To mechanistically address whether early-life LCS consumption impacts memory function, here we developed a model in which juvenile rats (postnatal [PN] day 26) are given daily access to LCS (saccharin, stevia, or acesulfame potassium [ACE-K]) under conditions where oral consumption is voluntary, the daily dose is fixed based on body weight and is within the U.S. FDA-recommended ADI. Our goal was to investigate the impact of voluntary daily LCS consumption throughout the juvenile and adolescent stages of development in rats (PN 26-60) on ingestive and cognitive behavioral outcomes during adulthood, and whether early life LCS consumption leads to changes in glucoregulatory function, the gut microbiome, and neuronal gene expression patterns in adulthood.

## 2. Methods

### 2.1 Animal Monitoring

Male and female Sprague Dawley rats (Envigo, Indianapolis, IN, USA; postnatal day (PN) 25; 50-70g) were housed individually in a climate controlled (22–24 °C) environment with a 12:12 light/dark cycle (lights off at 6pm). Rats were maintained on standard chow (Lab Diet 5001; PMI Nutrition International, Brentwood, MO, USA; 29.8% kcal from protein, 13.4% kcal from fat, 56.7% kcal from carbohydrate) and water. All experiments were performed during the light cycle. At PN 26, rats were randomized into groups of comparable weights and provided with their experimental diets. Body weights were measured daily whereas water consumption and chow intake were measured 3 times per week. All experiments were approved by the Animal Care and Use Committee at the University of Southern California and performed in accordance with the National Research Council Guide for the Care and Use of Laboratory Animals.

### 2.2 Experiment 1

#### Diet

Juvenile male and female rats (n=10 per sex/sweetener) were provided with the maximum acceptable daily intake (ADI) in mg/kg body weight, as recommended by the U.S. FDA, for acesulfame potassium (ACE-K; catalog # A2815, Spectrum Chemical, Gardena, CA, USA; 0.1% weight/volume (w/v) in reverse osmosis (RO) water; ~15 mg/kg), saccharin (catalog # 81-07-2, Sigma, St. Louis, MO, USA; 0.1% w/v in RO water; ~15 mg/kg), or stevia (JG Group, Ontario, Canada; 0.033% w/v in RO water; ~4 mg/kg) from PN 26-77 (approximately 7 weeks of access to the assigned sweetener). The volume required for delivery of each solution was calculated based on body weight daily and injected into empty rodent sipper tubes, fastened at the end with vinyl caps (6mm inner diameter, Jocon SF9000) to prevent leakage, and placed on the wire rack of the home cage adjacent to the rat’s *ad libitum* standard chow and water bottle. Voluntary consumption of the entire sweetener ration was verified daily by inspecting the tube for all animals. Rats in the control group (CTL; n=10 per sex) were provided a sipper tube filled with RO water at an equivalent volume/body weight as the ACE-K and saccharin groups. The concentration of each sweetener was selected based on in-house two bottle preference tests (concentrations preferred to water) as well as concentrations used in published studies (*31*, *32*). Sweetener access ceased at PN 77.

#### Behavioral Experiments

##### Novel Object in Context

Contextual episodic memory was assessed in rats beginning at PN 63 (following 30 days of LCS consumption) using the hippocampal-dependent Novel Object in Context (NOIC) task, a timeframe and behavioral procedure adapted from (*33*). The NOIC procedure took place over 5-days, each day consisting of a 5-min session per animal. The apparatus and objects were cleaned with 10% ethanol (EtOH) between each animal. On days 1 and 2, rats were placed in Context 1, a semi-transparent box (38.1 cm W × 61 cm L × 30.5 cm H) with yellow stripes, or Context 2, a grey opaque box (43.2 cm W × 43.2 cm L × 40.6 cm H) (one context/day in counterbalanced order). Following this habituation phase, each animal was placed in Context 1 containing Object A and Object B placed on diagonal, equidistant markings with ample space for the rat to circle the objects (NOIC, Day 1). Object A and Object B were an unopened 12 oz. coke can and a stemless wine glass, counterbalanced among animals. Importantly, the side Object A was located on was counterbalanced by group. The following day (NOIC day 2), rats were placed in Context 2 with identical copies of Object A. On the test day (NOIC day 3), rats were placed again in Context 2, except this time with Object A and Object B (which was not a novel object per se, but its placement in Context 2 was novel to the rat).

On NOIC days 1 and 3, exploration (defined as sniffing or touching the object with the nose or forepaws) was hand-scored by an experimenter blinded to the experimental group assignments who was viewing a live camera recording displayed on a computer monitor. The discrimination index for Object B [Time spent exploring Object B/ (Time spent exploring Object A + Time spent exploring Object B)] was calculated for days 1 and 3. Data were represented as % shift from baseline, where baseline is the discrimination index on day 1.

##### Barnes Maze

To test for hippocampal-dependent spatial memory, we employed a Barnes Maze task as previously described (*34*, *35*). In this task, rats were placed on a Barnes Maze (Med Associates; St Albans, VT, USA), a circular elevated platform (Diameter: 122 cm, Height: 140 cm) containing 18 identical holes spaced twenty degrees apart along the edge. Four sets of visuospatial cues (e.g., black and white stripes, a white circle, a stuffed unicorn, and an assortment of irregular shapes) were displayed on the room walls surrounding the maze, approximately 1 meter from the edges of the maze. The rats were habituated to the maze for 1 day as previously described (*35*), then trained for two days with two trials per day (as described below). The probe test, which assesses spatial memory retention, was conducted at PN 77.

During each training trial, the rat was placed in a start box for 30 seconds, then the box was lifted and the animal was given 3 minutes to find the hidden escape box within one of the holes. Each rat was assigned a specific escape hole in relation to the spatial cues, with the location counterbalanced across groups. To motivate the rats to search for the escape box, mildly aversive stimuli (120 W bright overhead light and 75 db white noise) were used, with the white noise being shut off when the rodent entered the escape box (*34*). After each trial, the rat was left undisturbed in the escape box for 1 minute before being returned to its home cage. Between each rat and trial, all surfaces were cleaned with 10% EtOH and the maze was rotated 180 degrees (to eliminate olfactory strategies). In the case that the rat failed to find the escape hole within the 3-minute trial, the experimenter placed the rat inside the escape hole for 1 minute. Latency (defined as the time to reach the escape hole), and the number of incorrect hole investigations was recorded by the experimenter. The procedure for the probe test was similar to training, except that the test was a single 2-minute trial with no escape box. Data were presented as the percent correct investigations on the first 10 investigated holes.

##### Zero Maze

All rats were tested for anxiety-like behavior in the Zero Maze on PN 110 or 111. The Zero Maze was an elevated circular platform (63.5 cm height, 116.8 cm external diameter) with two closed zones and two open zones, all of which were equal in length. The closed zones were enclosed with 17.5 cm high walls whereas the open zones had only 3 cm high curbs. Animals were started in the same closed arm of the maze and allowed to roam the maze for a single 5-minute session. After each session, the apparatus was cleaned with 10% EtOH. An experimenter scored the total time the rat spent in the open zones. A rat was considered in an open zone if its head and two front paws were in the open zone, as previously described (*35*). Data were reported as percent time spent in the open zones over the 5-minute test.

#### Ingestive behavior

##### Solutions

Corn oil emulsions were made by mixing 4.5% oil (Mazola, ACH Foods Inc., Oakbrook Terrace, IL, USA) and 0.6% of an emulsifier (Emplex®) in deionized water (dH_2_O) in an emulsifying blender before the training sessions. Emulsions were blended again if oil droplets started to appear in the solution. Reagent-grade glucose (0.56 M), fructose (0.56 M), 10% w/v MALTRIN (Maltrin580), quinine HCl (0.15 mM, 0.3 mM, and 1mM; QHCl), and lithium chloride (0.12 M LiCl) were prepared fresh with dH_2_O before each lickometer session. While the concentration of LiCl produces aversion, consumption of LiCl was capped to prevent over-ingestion, based on (*36*).

##### Lickometer training and tests

Animals were given 30-minute sessions (with the exception of the LiCl test, which was 20 minutes, see below) in identical operant chambers equipped with optical lickometers (Habitest, Coulbourn Instruments, Allentown, PA, USA). The optical lickometer registered licks from a sipper spout via breaks in a photobeam positioned just in front of the spout orifice. The sipper spout was in the recessed magazine in the center of one end wall, ~2 cm above a grid floor. Access to the sipper spout was computer-controlled via a motorized guillotine door. Licks were timestamped and recorded via Coulbourn Instruments’ Graphic State software (Ver 4.0).

Starting at PN 82, overnight water deprived rats were trained to lick in the lickometer for 30-minute sessions on two consecutive days (one session / day). On each day, rats were offered a bottle of dH_2_O in the lickometer. The second day, any rat that did not take at least 800 licks in 30 minutes was given an additional 30-minute session with dH_2_O later the same day. After the second training day, rats were provided home cage water to replete. Then, starting on PN 85, rats were provided rations every day to maintain their body weight to 85% of their *ad libitum* feeding body weight. At PN 88, rats were trained to lick for 4.5% corn oil emulsion for one 30-minute session so they would associate the spout with calorie intake. Rats were retrained as necessary the same day, under the same criterion as water training (at least 800 licks at the end of the session). Testing started at PN 89 and occurred over two days. Testing order was counterbalanced such that half the animals were given 0.56 M glucose to consume for 30 minutes on the first test day and 0.56 M fructose on the second test day; the other half received the reverse order. After one day of rest, rats were then tested with 10% w/v MALTRIN for 30 minutes and were returned to *ad libitum* chow 30 minutes after testing. From PN 95-105, the rats were water restricted (bottles pulled the day before water retraining, which occurred over 1 day, and returned 30 minutes after the test) to test for 30-minute quinine intake (starting with the lowest concentration on PN 96 and ending with the highest concentration on PN 105, with 2-4 days on *ad libitum* water between concentrations to allow for sufficient rehydration). At PN 130, overnight water deprived rats were retrained with 30-minute access to water, and then water deprived again. The next day, rats were tested with the 0.12 M LiCl solution. Microstructural analyses of licking patterns on each test were performed with the time stamped lick records using an open-source Python program (https://github.com/pungaliy/Microstructural_Lick_Analysis). Bursts were defined as runs of licks, separated by a ≥ 1 second pause in licking (*37*). In addition to the total number of licks taken per session, total licks in the first minute, first burst size, number of bursts, and mean burst sizes were computed.

#### Progressive ratio operant responding for sucrose or high-fat diet

Operant response training for sucrose was conducted as previously described (*38*). Rats were habituated to 20 sucrose pellets in the home cage on PN 134. Starting at PN 135, the rats received a 1-h training session each day over 6 days in standard conditioning boxes (Med Associates, Fairfax, VT, USA) that contain an ‘active’ lever and an ‘inactive’ lever, whereby pressing the active lever, which was the same for each animal, results in the release of a 45 mg sucrose pellet into a food cup (F0023, Bio-Serv, Frenchtown, NJ, USA). The first two days, the rats were trained to press the active lever on a fixed ratio-1 (FR1) schedule with an autoshaping component. On these sessions, each active lever press results in sucrose reinforcement. In the case that 10 minutes lapse without an active lever press, a pellet is automatically dispensed. The next four days of training consisted of FR1 followed by FR3 (3 active lever presses were required to obtain 1 pellet) training without autoshaping for 2 days each. Subsequently, from PN 151-152, the rats were tested using a progressive ratio (PR) reinforcement schedule. As previously described, the number of lever presses for a sucrose pellet increased progressively using the following formula:

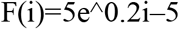

where F(i) is the number of lever presses required for the next pellet at i=pellet number (*38*, *39*). The test session terminates when the rat did not achieve a ratio within 20 minutes, and the total number of pellets earned was recorded. Beginning on PN 153, rats were retrained with two FR1 sessions and one FR3 session (one session / day) with high-fat pellets (35% kcal fat enriched with sucrose, F05989, Bio-Serv, Frenchtown, NJ, USA). Then, the rats were retested from PN 159-160 on a PR schedule, as above, except that high-fat pellets were used in place of sucrose pellets.

#### Free access sucrose consumption in the home cage

From PN 165-193, all rats were provided *ad libitum* access to an 11% w/v sucrose solution (C&H Pure Granulated White Cane Sugar, Crockett, CA, USA; dissolved in RO water) in addition to standard chow and a water bottle in the home cage. The concentration of sucrose was selected to match the one found in common sugar-sweetened beverages consumed by humans (*40*) and our prior studies (*35*). Sugar intake was measured and fresh sugar solutions were provided every three days. After 4 weeks of access, the average sucrose consumed per day (in kcals) in sweetener groups was compared to intake of sucrose in CTL groups.

#### Body Composition

At PN 194, rats were food restricted for one hour prior to being weighed and scanned for body composition as previously described (*7*) using the Bruker NMR Minispec LF 90II (Bruker Daltonics, Inc., Billerica, MA, USA). The apparatus was based on Time Domain Nuclear Magnetic Resonance (NMR) signals from all protons and has the benefit of being non-invasive and requiring no anesthesia. Percent body fat was calculated as [fat mass (g)/body weight (g)] x 100.

#### Microbiome

##### Fecal collection

Immediately after Barnes Maze testing (PN 77), animals were individually placed in a sterile cage with no bedding and were mildly restrained until defecation occurred. Fecal samples were weighed under sterile conditions and then placed into a DNAse/RNAse free 2 mL cryogenic vial embedded in dry ice. Samples were then stored in a −80°C freezer. All materials used to collect samples were cleaned with 70% EtOH in between rats.

##### 16s rRNA Sequencing

Frozen fecal samples were shipped on dry ice to UC Davis MMPC and Host Microbe Systems Biology Core. Total DNA was extracted using Mo-Bio (now Qiagen) PowerFecal kit. Sample libraries were prepared and analyzed by barcoded amplicon sequencing. In brief, the purified DNA was amplified on the V4 region of the 16S rRNA genes via PCR using the following primers: F319 (5’-ACTCCTACGGGAGGCAGCAGT-3’) and R806 (5’-GGACTACNVGGGTWTCTAAT-3’). High-throughput sequencing was performed with Illumina MiSeq paired end 250-bp run.

##### Sequence processing and data analysis

Sequencing reads were analyzed with DADA2 and QIIME2 following the developers’ instructions (*41*). The forward reads were truncated to 200bp and denoised to amplicon sequence variants (ASVs), and the chimera sequences were detected with the DADA2 ‘consensus’ method and removed. The ASV sequences were classified using the QIIME2 sklearn classifier with the SILVA database (release 132) (*42*). The taxonomic abundance tables were normalized as previously described to correct for the varied sequencing depth across samples (*43*). The statistical analysis was performed with R. The PCoA ordinations of the microbiomes were calculated based on the Bray-Curtis dissimilarity and visualized using the R function ‘capscale’ in the package ‘vegan’. The PERMANOVA test of the associations between microbiome and groups (CTL vs LCS, and CTL vs ACE-K) was performed with the function ‘adonis’ in the same package. Shannon diversity index was calculated with the function ‘diversity’ and used to characterize the alpha diversity of the microbiomes. The associations of individual taxa and groups (CTL vs LCS, and CTL vs ACE-K) were analyzed with a linear regression model with Group and Sex of the rat as the main effects and Group x Sex as the interaction. Rare taxa (prevalence <25% samples) were not included in order to avoid over adjustment for false discovery rate. The Wilcoxon test (FDR<0.1) was used to identify significant differential taxa between treatment and CTL. The P-values were adjusted for multiple hypotheses testing with the Benjamini-Hochberg method.

### 2.3 Experiment 2

To examine the effects of early life sugar and LCS consumption on glucose tolerance in adulthood, a cohort of juvenile male rats (n=15) was divided into the following two groups at PN 26: 1) control diet, with standard chow and water, and a sipper tube injected daily with water as in Experiment 1 (CON; n=8), 2) ACE-K diet, which received the U.S. FDA-recommended ADI amount of ACE-K (0.1% w/v; ~15mg/kg) daily via a sipper tube, with standard chow and water as described above (ACE-K; n=7). Rats were on the experimental diets until PN 74, to approximately match the ACE-K exposure in Experiment 1 (7 weeks of access), and then subsequently outfitted with gastric catheters from PN 79-81 to compare glucose tolerance following either oral or gastric dextrose administration.

#### Gastric catheter surgeries

Gastric catheter surgeries were conducted as described in Schier *et al*. previously (*44*). Briefly, following an overnight fast, rats were laparotomized while under the anesthetic isoflurane (~5% induction rate; ~1.5–3% maintenance rate). A gastric catheter made of silastic tubing (inside diameter = 0.64 mm, outside diameter = 1.19 mm; Dow Corning, Midland, MI, USA) was inserted ~1 cm into the stomach through a puncture wound in the greater curvature of the forestomach. The catheter was tethered to the stomach wall with a single stay suture, silastic cuff, and piece of Marlex mesh (Davol, Cranston, RI, USA). A purse string suture and concentric serosal tunnel were used to close the wound in the stomach. The other end of the catheter was then tunneled subcutaneously to an interscapular exit site, where it was attached using a single stay suture and a larger square piece of Marlex mesh. The tube was then connected to a Luer lock adapter, as part of a backpack harness worn by the rat (Quick Connect Harness, Strategic Applications, Lake Villa, IL, USA). Rats were treated postoperatively with gentamicin (8 mg/kg, subcutaneous injection) and ketoprofen (1 mg/kg, subcutaneous injection). Rats were given increasing increments of chow (1-3 pellets) after surgery and then *ad libitum* access to chow. The gastric catheter was routinely flushed with 0.5 ml of isotonic saline beginning 48 h after surgery to maintain its patency. Harness bands were adjusted daily to accommodate changes in body mass.

#### Intragastic and Oral Glucose Tolerance Test

The method for comparing the effects of oral versus intragastric (IG) glucose tolerance tests in rats was newly established and modified from prior procedures (*27*, *45*). Following recovery from surgery, all rats were trained to lick at the lickometer for water and habituated to passive IG infusions in the Coulbourn chambers. After an overnight water deprivation, rats were given 30-minute free access to a bottle and sipper containing water, and then returned to the home cage (without water). Approximately 3-4 hours later, rats were returned to the chambers. The IG catheter was connected to an infusion line, which consists of polyethylene tubing encased in a spring tether and routed through a single channel swivel allowing the rats to move freely about the chamber. The other end of the infusion line was connected to syringe in a computer-controlled infusion pump. Rats were infused with 4 ml of water at a rate of 0.75 ml/minute. Rats were returned to the home cage at the end of the IG infusion; no home cage water was provided. On the following day, rats received the same IG habituation session in the morning. 3-4 hours later, rats received a second lickometer training session. In this session, rats had access to water for 5 minutes through a sipper spout that was connected to an infusion pump. In this system, licking the spout activated the infusion pump for 1 seconds at a rate of 0.75 ml/minute and during this activation time licks did not re-activate the pump. Rats were returned to a home cage with water after this session. Then, to train the rats to associate the spout with nutritive value, a third training session was conducted with 4.5% w/v corn oil emulsion in place of water, for 30 minutes, in the morning following a 20 hour fast. Animals were given additional 30-minute training sessions if they did not take at least 800 licks. Approximately 3-4 hours later, rats were placed back in the chamber and received a 4 ml IG infusion of corn oil over 5 minutes. Chow was returned after the IG infusion.

All rats underwent oral and IG glucose tolerance testing in counterbalanced order on PN 98 and PN 101. Chow was removed from the home cage 20 hours prior to test. Water was removed from the home cage approximately 14-20 hours prior to testing to encourage licking during the test. Then, five minutes prior to the start of each test, baseline blood glucose readings were collected from the tip of the tail and measured using a glucometer (One touch Ultra2, LifeScan Inc., Milpitas, CA, USA). At minute 0, rats received 3 ml of a 20% w/v dextrose solution dissolved in sterile saline via a pump-infused licking session (OGTT) or IG infusion (IGTT) over 5 minutes. Blood glucose readings were obtained at 5, 10, 30, 60, and 120 minutes after time 0. Rats were kept in the behavioral chambers until after the 30-minute blood glucose measurements and were promptly returned to their home cages for the remaining measurements.

### 2.4 Experiment 3

Male and female rats (n=16 per sex/treatment) were given daily ACE-K (0.1% w/v; ~15mg/kg) through a sipper tube, chow, and water (ACE-K group); or chow, water, and a sipper tube with water (CTL group) until PN 80. Our goal was to match the design of Experiment 1 (~7 weeks access to ACE-K), in order to generate tissues for analyses. At PN 80, chow and ACE-K sippers were removed 4 hours into the light cycle and all rats were anesthetized with a Ketamine:Xylazine:Acepromazine mixture (90mg/kg: 2.8mg/kg: 0.72mg/kg, intramuscular injection) before being rapidly decapitated for collection of circumvallate tissue in RNAlater (ThermoFisher Scientific, Waltham, MA) and flash frozen brain tissue.

#### qPCR

##### Circumvallate tissue harvest

Given that sugar-sensing genes can display a diurnal rhythm (*46*) and gene expression within the circumvallate (CV) has a circadian rhythm (*47*), CV samples analyzed were collected within a two-hour timeframe six hours after fasting. The whole tongue was removed and pinned into a Sylgard® dish filled with a Tyrode’s solution (140 mM NaCl, 5 mM KCl, 2 mM CaCl_2_, 1 mM MgCl_2_, 10 mM HEPES and 10 mM glucose). Under a dissecting microscope, 3 ml of an enzyme cocktail [1 mg/ml collagenase A (#11088793001, Sigma-Aldrich, St. Louis, MO, USA) and 0.1 mg/ml elastase (Sigma-Aldrich, St. Louis, MO, USA) in phosphate-buffered saline (PBS)] was injected under the tongue epithelium, and the tongue was incubated at 37°C for approximately 20 minutes. Then, the epithelium was carefully peeled from the underlying connective tissue under the dissecting microscope, and the CV was separated from the rest of the epithelium. The CVs were stored overnight at 4°C in RNAlater (ThermoFisher Scientific, Waltham, MA), then transferred in an empty tube and stored at −80°C.

##### Reverse Transcriptase-quantitative Polymerase Chain Reaction (RT-qPCR)

To quantify the relative *Tas1r2* and *Tas1r3* mRNA expression between the control (n= 8, 4 females and 4 males) and ACE-K (n= 8, 4 females and 4 males) rats in the CV part of the tongue epithelium, reverse transcriptase quantitative polymerase chain reaction (RT-qPCR) was performed as previously described (*48*). Total RNA was extracted from each sample using the RNeasy Lipid Tissue Mini Kit (Cat No. 74804, Qiagen) following the manufacturer’s instructions. The total RNA concentration per sample was measured with a NanoDrop Spectrophotometer (ND-ONE-W, ThermoFisher Scientific). cDNA was synthesized using the QuantiTect Reverse Transcription Kit (Cat No. 205311, Qiagen) following the manufacturer’s instructions, and amplified using the TaqMan PreAmp Master Mix (Cat No. 4391128, ThermoFisher Scientific). Real-time PCR was performed using TaqMan Gene Expression Assay for rat β-actin (Actb, Rn00667869_m1, Applied Biosystems), rat Taste receptor type 1 member 3 (Tas1r3, Rn00590759_g1, Applied Biosystems) and rat Taste receptor, type 1, member 2 (Tas1r2, Rn01515494_m1, Applied Biosystems), and TaqMan Fast Advanced Master Mix (Cat No 4444557, Applied Biosystems), in the Applied Biosystems QuantStudio 5 Real-Time PCR System (ThermoFisher Scientific). All reactions were run in triplicate, and controls wells without cDNA template were included to verify the absence of genomic DNA contamination. The triplicate Ct values for each sample were averaged and normalized to β-actin expression. The comparative 2-ΔΔCt method was used to quantify the relative expression levels of our genes of interest between groups.

#### RNA-seq

##### Brain tissue collection (dHPC and ACB)

Whole brains were removed, flash frozen in isopentane surrounded by dry ice and stored at −80°C until use. Tissue punches of dorsal hippocampus (dHPC) and nucleus accumbens (ACB) (2.0 mm circumference, 1-2 mm depth) were collected from brains mounted on a stage in a Leica CM 1860 cryostat (Wetzlar, Germany). Anatomical landmarks based on the Swanson brain atlas [dHPC containing dorsal cornu ammonis area 1 and dorsal dentate gyrus at atlas levels 28-30, and ACB at atlas levels 10-12)] (*49*).

##### Sample preparation and sequencing

Total RNA was extracted according to the manufacturer’s instructions using an AllPrep DNA/RNA Mini Kit (Qiagen, Hilden, Germany) and checked for purity using the NanoDrop One (Thermo Fisher Scientific, Waltham, MA, USA). All samples were found to have high purity and were sent to the USC Genome Core for library preparation and RNA sequencing. There, the total RNA was checked for degradation using a Bioanalyzer 2100 (Agilent, Santa Clara, CA, USA) to verify the quality for all samples. Libraries were prepared from 1 ug of total RNA using a NuGen Universal Plus mRNA-seq Library Prep Kit (Tecan Genomics Inc. Redwood City, CA, USA). Final library products were quantified using the Qubit 2.0 Fluorometer (Thermo Fisher Scientific Inc., Waltham, MA, USA), and the fragment size distribution was determined with the Bioanalyzer 2100. The libraries were pooled equimolarly, and the final pool was quantified via qPCR using the Kapa Biosystems Library Quantification Kit, according to manufacturer’s instructions. The pool was sequenced using an Illumina NextSeq 550 platform (Illumina, San Diego, CA, USA), in Single-Read 75 cycles format, obtaining about 25 million reads per sample.

##### RNA-sequencing quality control

RNA-seq quality control was performed using FastQC (*50*). Low-quality reads were trimmed by Trimmomatic (*51*). Reads were then aligned to Rattus norvegicus genome Rnor6.0 using STAR (*52*). Gene counts were quantified using HTSeq (*53*). Principal component analysis (PCA) was used to detect potential sample outliers, and one ACB sample from the ACE-K treatment group was removed.

RNA-seq quality control was performed using FastQC (*50*). Low-quality reads were trimmed by Trimmomatic (*51*). Reads were then aligned to Rattus norvegicus genome Rnor6.0 using STAR (*52*). Gene counts were quantified using HTSeq (*53*). PCA was used to detect potential sample outliers (one ACB sample from the ACE-K treatment group was removed).

##### Identification of differentially expressed genes (DEGs)

Genes detected in less than three samples or with a normalized count lower than five were filtered out. DEseq2 was then used to perform differential gene expression analysis between control and ACE-K treatment groups across both sexes to identify DEGs affected by ACE-K Treatment, Sex, and Treatment x Sex interactions, or within males and females separately to identify DEGs affected by treatment within each sex (*54*). P-values were adjusted for multiple testing corrections using Benjamini-Hochberg correction. At false discovery rate (FDR) < 0.05, no significant DEGs were identified for the treatment effect in both across and within sexes analyses. Suggestive DEGs were defined as DEGs with an unadjusted P-value < 0.05 and the absolute value of log fold change >= 0.4. For heatmap visualization, raw counts were normalized using regularized log transformation implemented in DESeq2. Z-scores for each gene were then calculated and visualized.

##### Pathway analyses of DEGs

Suggestive DEGs were selected for pathway analyses. Pathway analysis was conducted using EnrichR by checking the DEG enrichment in curated pathways from KEGG, BioCarta, Reactome (https://reactome.org/), and gene ontology (GO) biological pathways (*55*–*57*). Pathways with an FDR < 0.05 were considered significant.

### 2.5 Statistics

All data except for gene and microbiome sequencing results are presented as mean ± SEM and analyzed and graphed using Prism software (GraphPad Inc., Version 9.1.2 (225)) or Statistica (Version 7; Statsoft), with significance set as p < 0.05. Body weights, water and caloric intake were analyzed using a multi-factor mixed ANOVA with Time as a within-subjects factor and Group (Experiments 1-3), Sex (Experiments 1,3) and Sweetener (Experiment 1) as between-subjects factors. Glucose tolerance results were analyzed via 2-way mixed ANOVA with Time as a within-subjects variable and Group as a between-subjects variable. Body composition, NOIC, Barnes Maze, Zero Maze, licking/ingestive tests, progressive ratio, sucrose consumption in the home cage, and *Tas1r2* and *Tas1r3* mRNA expression were analyzed using a multi-factor ANOVA with Sex, Group, and Sweetener as the independent between-subjects variables (except for *Tas1r2* and *Tas1r3* mRNA analyses which did not include Sweetener as a variable). Data were corrected for multiple comparisons using Sidak’s multiple comparison test. Given that there was not a significant interaction or main effect of Sex or Sweetener in the majority of analyses from Experiment 1 (exceptions being Barnes Maze and the glucose vs. fructose test), all of the data from Experiment 1 are also presented with sexes and sweeteners combined as the “LCS” group, as well as separated.

## 3. Results

### 3.1 Early life LCS consumption impairs post-oral glucose tolerance without influencing body weight, caloric intake, water intake, or adiposity

Early life LCS consumption did not yield differences in body weight (Fig. 1a, Supplemental Fig. 2a-h), caloric intake (Fig. 1b, Supplemental Fig. 2i-p), water intake (Fig. 1c), or body composition (Supplemental Fig. 2q) relative to controls. Despite the lack of group differences in these energy balance parameters, early life LCS-fed (ACE-K, specifically) rats showed impaired post-oral glucose tolerance, as evident from higher blood glucose levels at 30 minutes after intragastric administration of a glucose load that bypasses the oral cavity (Fig. 1d) (*P* = 0.0244 for Group). On the other hand, LCS-exposure was not associated with differences in blood glucose clearance when the same glucose load was consumed orally (Fig. 1e).

**Figure 1.**
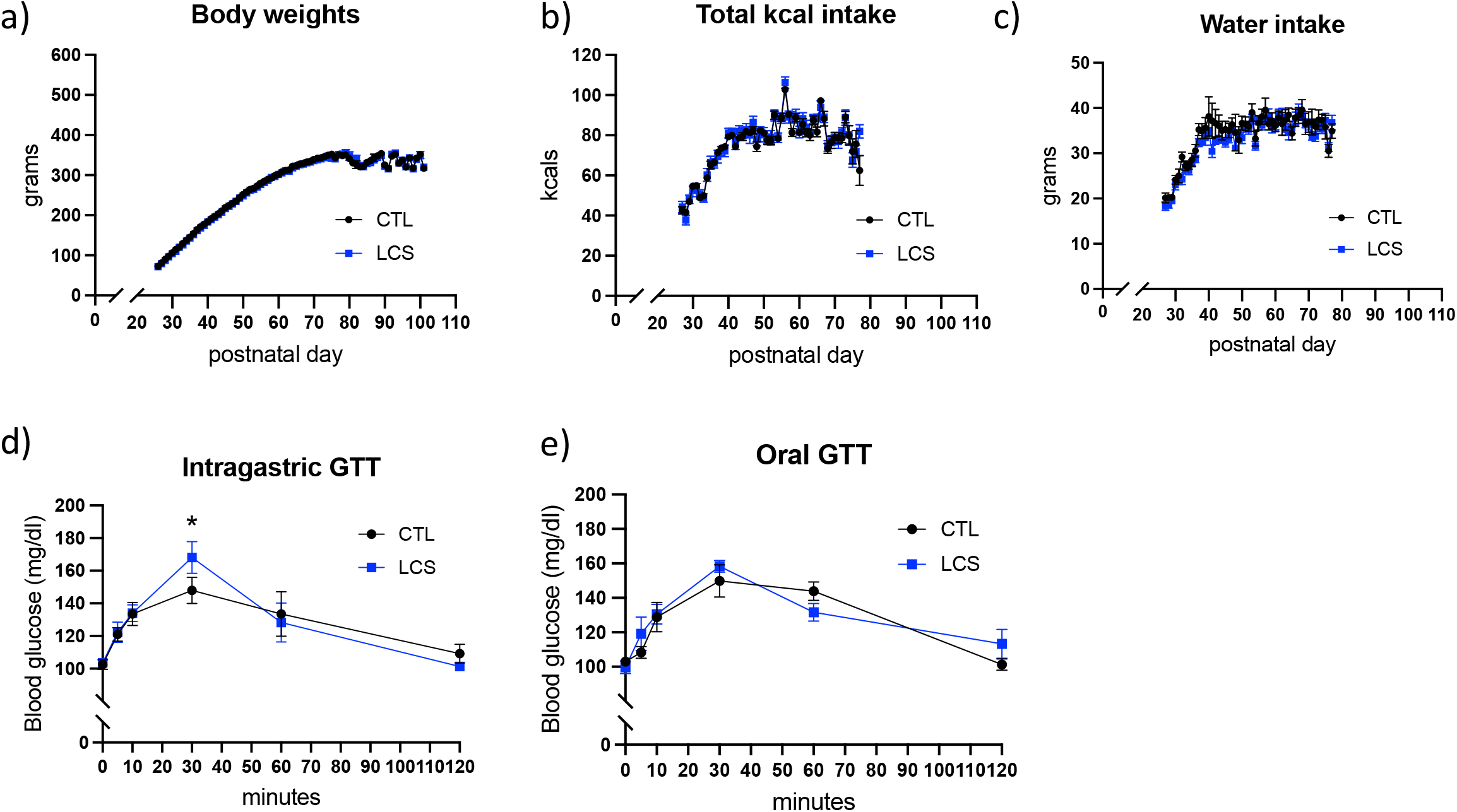
Early life LCS consumption impairs peripheral glucose metabolism without influencing total caloric intake, body weight, or adiposity. Daily LCS consumption (acesulfame-K, saccharin, stevia) during development did not affect body weight (a), total caloric intake (b), or water intake (c). Glucose intolerance in LCS-exposed rats was observed following intragastric administration of glucose (d), but not following oral glucose consumption when cephalic phase and orosensory responses to glucose were intact (e). Data are means ± SEM; *P < 0.05, CTL: control; GTT: glucose tolerance test; kcals: kilocalories; LCS: low-calorie sweeteners

### 3.2 Early life LCS consumption impairs hippocampal-dependent memory in adulthood

In the NOIC task (depicted in Fig. 2a), rats normally spend more time exploring the object that is novel to the test context; this phenomenon is disrupted by hippocampal loss of function (*58*). Present results show that daily LCS consumption during the juvenile and adolescent period yielded contextual episodic memory impairments in the hippocampal-dependent NOIC task (timeline of experiment depicted in Supplemental Fig. 1a), exhibited by a lower shift from baseline discrimination index for investigation of the novel object in the test context for combined male and female LCS groups relative to controls. This outcome did not differ by sex or sweetener (sexes and sweeteners combined analysis in Fig. 2b [*P* = 0.0341 for Group], separated by sex in Fig. 2c, separated by sex and sweetener in Supplemental Fig. 3a-b). Spatial memory deficits associated with early life LCS consumption were also observed in the hippocampal-dependent Barnes Maze task (depicted in Fig. 2d), albeit only in the males, as demonstrated by significantly fewer correct (or adjacent to correct) hole investigations during the memory probe. This outcome did not differ by sweetener and was observed in males only (sexes and sweeteners combined in Fig. 2e, separated by sex in Fig. 2f [*P* = 0.0166 for Group in males], separated by sex and sweetener in Supplemental Fig. 3c-d). For females, neither control nor LCS animals utilized a spatial strategy to solve the Barnes maze as indicated by memory performance that was not above the probability due to chance (chance performance represented by the dashed line in Fig. 2f; 1-sample t tests vs. chance performance [0.167%] not significant for either group).

**Figure 2.**
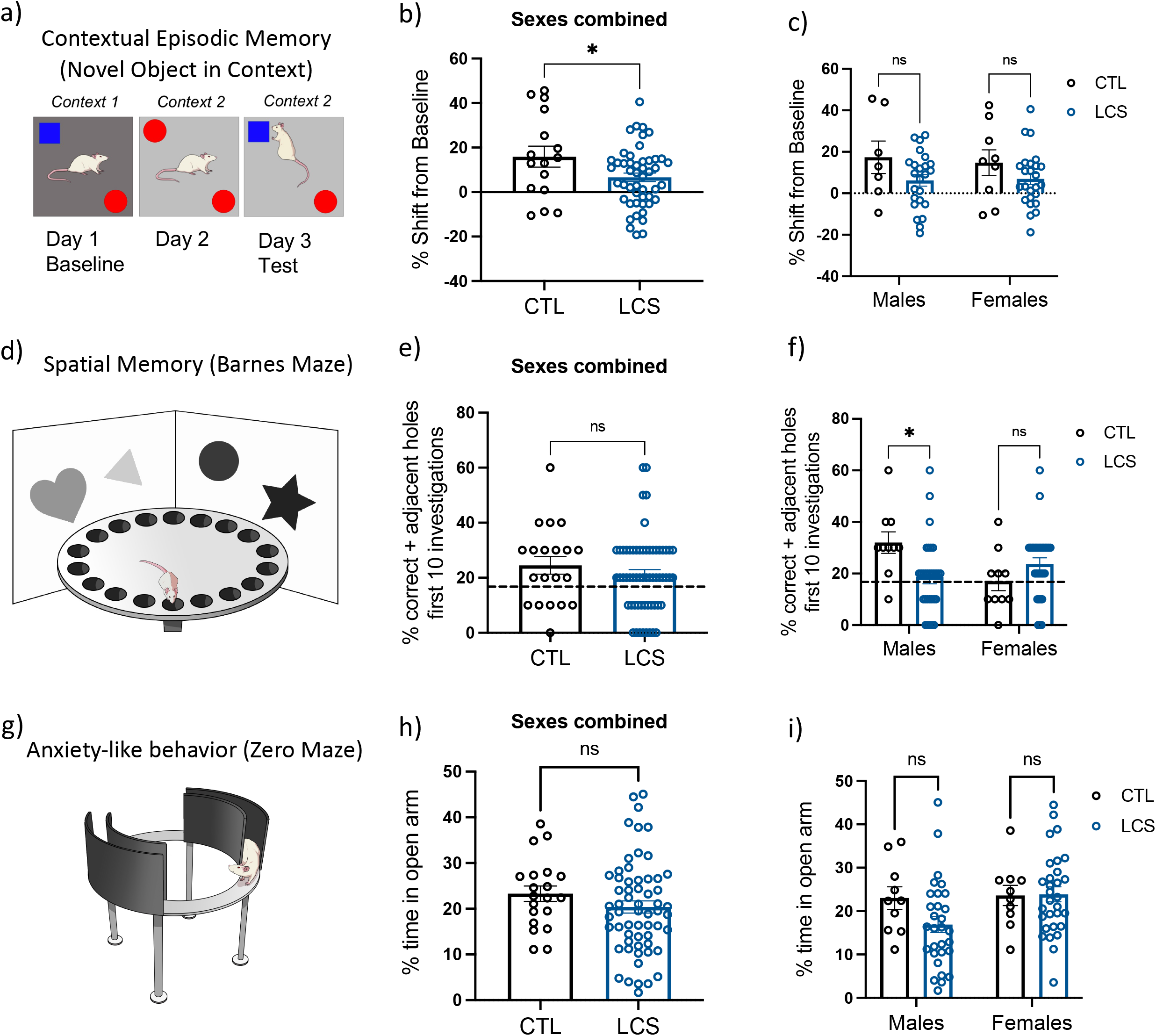
Early life LCS consumption impairs hippocampal-dependent memory during adulthood. LCS consumption (acesulfame-K, saccharin, stevia) during the juvenile and adolescent developmental stages impaired contextual episodic memory in the NOIC procedure (a) regardless of sex (b, c). LCS-associated spatial memory deficits in the Barnes Maze procedure (d) were observed in males, whereas neither female CTL nor LCS-exposed rats utilized a spatial strategy (e, f). There were no group or sex differences in anxiety-like behavior in the Zero Maze procedure (g-i). Data are means ± SEM; ns = nonsignificant, *P < 0.05, CTL: control; LCS: low-calorie sweeteners; NOIC: novel object in context

Early life LCS was not associated with significant group differences in anxiety-like behavior in the Zero Maze test (procedure depicted in Fig. 2g; percentages of time in the open arm area data are depicted in Fig. 2h [sexes and sweeteners combined], Fig. 2i [separated by sex], and Supplemental Fig 3e-f [separated by sex and sweetener]).

### 3.3 Early life LCS consumption alters ingestive responses to sugar and reduces lingual sweet taste receptor expression (Tas1r2 and Tas1r3) without affecting ingestion of bitter, salty, or non-sweet carbohydrate substances

Rats that consumed LCS daily during the juvenile and adolescent period demonstrated altered short-term sugar intake in adulthood (procedure depicted in Fig 3a). This was due, in part, to differential early taste-guided licking responses to the two sugars in the first minute of the test, when consummatory behaviors are primarily driven by taste and other cephalic cues. No such differences in early licking responses were observed in the controls. Post-hoc analyses revealed that, independent of sex and sweetener, LCS rats consumed more fructose during the first minute of consumption relative to equimolar glucose (sexes and sweeteners combined in Fig. 3b [*P* = 0.0002 for Sugar], separated by sex in Fig. 3c-d [*P* = 0.0238 for Sugar in males, *P* = 0.0083 for Sugar in females], separated by sex and sweetener in Supplemental Fig 4a-b). Across the entire session, however, control rats licked more for glucose than fructose. This is consistent with previous studies showing that glucose stimulates ingestion from a post-oral site of action more so than fructose (*59*). By contrast, LCS rats consumed comparable amounts of the two sugars in the 30-minute test (Fig. 3e; *P* = 0.3654), an outcome primarily driven by males with a significant Group x Sugar interaction (*F*_(1, 32)_ = 6.6981, *P* = 0.01440) and a significant main effect of Sugar (*P* < 0.001), but not Group. Post-hoc analyses revealed that there was a significant difference between glucose and fructose consumption in the control males (*P* < 0.001 Fig. 3f) but not in LCS males. In females, no significant interactions or main effects were observed (Fig. 3g). Monosaccharide consumption was not differentially influenced by individual sweetener as all main effects and interactions with sweetener were not significant (Supplemental Fig. 4c-d). There were no overall LCS-associated differences in intake of a non-sweet carbohydrate, MALTRIN, though ACE-K-exposed female rats licked significantly more for MALTRIN in the first minute of the test than did the females in the control group (Supplementary Fig 5a-h).

**Figure 3.**
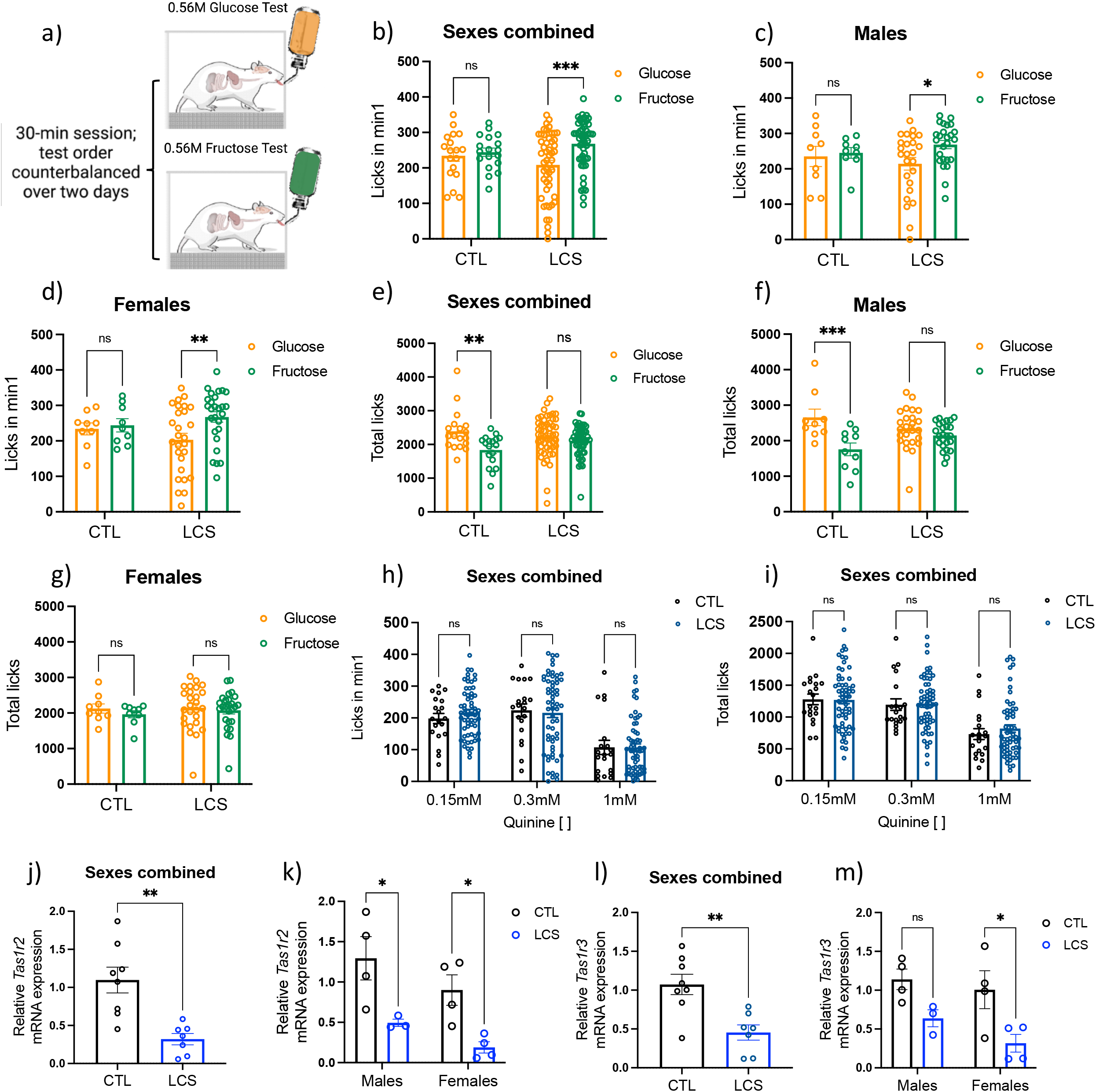
Early life LCS exposure alters sugar consumption and reduces lingual sweet taste receptor expression. Schematic displaying the use of the lickometer for analyses of ingestive responses to equimolar glucose and fructose solutions is depicted in (a). Whereas CTL showed equivalent short-term (first minute) ingestive responses for a glucose versus an equimolar fructose solution, rats previously given daily LCS treatment (acesulfame-K, saccharin, stevia) show heightened initial licking responses for the fructose solution relative to glucose regardless of sex (b-d). Longer-term (30 min) ingestive appetitive responses were higher for the glucose relative to the fructose solution in CTL, but not in LCS-exposed rats (e-g). Ingestive responding for the bitter tastant quinine, as measured by licks during the first minute of exposure (h), or whole-session consumption (i), was comparable between CTL and LCS-exposed animals. LCS-exposed rats (acesulfame-K) also had reduced gene expression levels of the sweet taste receptors, *Tas1r2* (j,k) and *Tas1r3* (l,m) in the CV regardless of sex. Data are means ± SEM; ns = not significant, *P < 0.05, **P < 0.01, ***P < 0.001. CTL: control; CV: circumvallate papillae of the tongue; LCS: low-calorie sweeteners

Although several LCSs have bitter and/or metallic taste qualities, LCS-exposed rats did not display differences in consumption of the prototypical bitterant, quinine, at various concentrations (0.15, 0.3, and 1 mM) during the first minute (Fig. 3h) or entire session (Fig. 3i). Intake of quinine separated by sex and by sweetener is shown in Supplemental Fig. 4e-t. While there were no overt differences in initial licking responses to a salt, lithium chloride, as a function of previous LCS exposure, LCS females consumed significantly less of it than control females (Supplemental Fig. 5i-o).

The altered ingestive responses to monosaccharides observed in rats given early life daily access to LCSs may be based on altered sweet taste receptor expression, as gene expression analyses for the two sweet receptors revealed reduced relative sweet taste receptor mRNA levels in the circumvallate papillae of LCS-exposed (ACE-K, specifically) rats compared to controls *[Tas1r2: P* = 0.0015; Fig. 3j, separated by sex in Fig. 3k, *P* = 0.0252 for males, *P* = 0.0311 for females) and *Tas1r3: P* = 0.0027; Fig. 3l, separated by sex in Fig. 3m, *P* = 0.0234 for females).

### 3.4 Early life LCS consumption reduces effort-based responses for sucrose yet increases free access sucrose consumption

Daily LCS consumption during the juvenile and adolescent period was associated with reduced motivation to lever press for sucrose, but not high fat, reinforcement in the operant progressive ratio task (depicted in Fig. 4a) when tested during adulthood (Group main effect for sucrose *P* = 0.0308; Fig. 4b), an effect primarily driven by the females (*P* = 0.0248 for females; Fig. 4c). In comparison, LCS-exposed rats earned similar amounts of HFD pellets during the progressive ratio test as controls (Fig. 4d, separated by sex in Fig. 4e).

**Figure 4.**
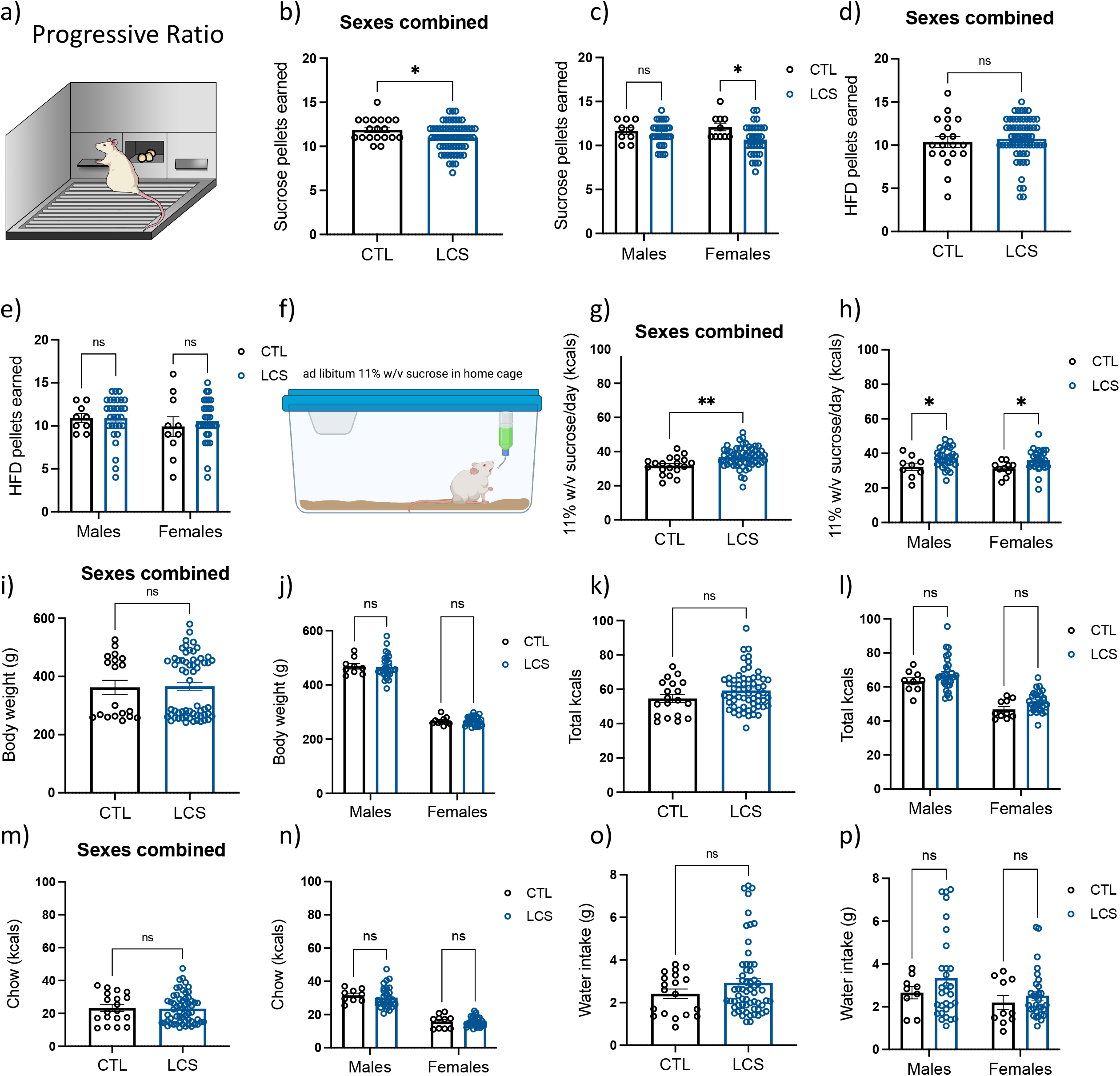
Early life LCS consumption reduces effort-based responding for sucrose while increasing long term free-access sucrose intake. In the progressive ratio schedule operant task which measures effort-based responding for food reinforcement (a), LCS rats (acesulfame-K, saccharin, stevia) earned fewer sucrose pellets collapsed across sex, though this effect was more pronounced in LCS females (b, c). No group differences were observed in motivated operant responding for high fat diet pellets (d, e). However, when provided with free access to a sucrose solution in the home cage (f), LCS rats consumed more of the sucrose solution relative to controls regardless of sex (g, h). There were no significant group differences in body weight (i, j), total (sucrose plus chow) caloric intake (k, l), caloric intake from chow (m, n), or water intake (o, p). Data are means ± SEM; ns = not significant, *P < 0.05, **P < 0.01. CTL: control; HFD: high fat diet; kcals: kilocalories; LCS: low-calorie sweeteners

In contrast, LCS-exposed rats showed increased free access consumption of sucrose (10% w/v) relative to controls when consumption was measured in the home cage over a 4-week period (Fig. 4f-g, *P* = 0.0011), an outcome that did not differ by sex (Fig. 4h, *P* = 0.0380 for males, *P* = 0.0443 for females). During these 4 weeks with sucrose access in the home cage, there were no long-term effects of early life LCS consumption on body weight (Fig. 4i, separated by sex in Fig. 4j), total calories consumed from chow and sucrose combined (Fig. 4k; separated by sex in Fig. 4l), calories consumed from chow (Fig. 4m; separated by sex in Fig. 4n), or water intake (Fig. 4o; separated by sex in Fig. 4p).

Neither the progressive ratio nor the home cage free access sucrose consumption results were significantly affected by sweetener (separated by sweetener in Supplemental Fig. 3g-t, no significant main effects or interaction with sweetener).

### 3.5 Collagen-related gene pathways in the hippocampus are altered by early life LCS consumption in a sex-specific manner

To explore potential mechanisms related to the effects of LCS consumption (ACE-K, specifically) on neurocognitive outcomes, brains from animals that received early life ACE-K were collected in adulthood for bulk RNA sequencing and gene pathway enrichment analyses (experimental timeline depicted in Supplemental Fig. 1c). Analysis of bulk rRNA sequencing of HPCd tissue punches (targeted region depicted in Fig. 5a) identified 132 differentially expressed genes (DEGs) for males and 138 DEGs for females (top 50 DEGs in HPCd-enriched tissue for each sex are depicted in Supplemental Fig. 6a-b). Despite the animals not having a clear group separation in the PCA, LCS consumption significantly altered gene pathways related to collagen formation and synthesis. Interestingly, various collagen-related pathways were all upregulated in LCS males relative to controls but were downregulated in LCS females relative to controls (Fig. 5b-d). Specifically, the DEGs were related to gene pathways involved in protein digestion and absorption (FDR = 0.0075, *P* = 3.74E-05 for males; FDR = 0.0075, *P* = 8.25E-05 for females), assembly of collagen fibrils and other multimeric structures (FDR = 0.0075, *P* = 3.51E-05 for males; FDR = 0.0006, *P* = 4.33E-06 for females), collagen biosynthesis and modifying enzymes (FDR = 0.01, *P* = 7.41E-05 for males; FDR = 0.0002, *P* = 6.69E-07 for females), and collagen formation (FDR = 0.0309, *P* = 0.0003 for males; FDR = 0.0006, *P* = 5.16E-06 for females).

**Figure 5.**
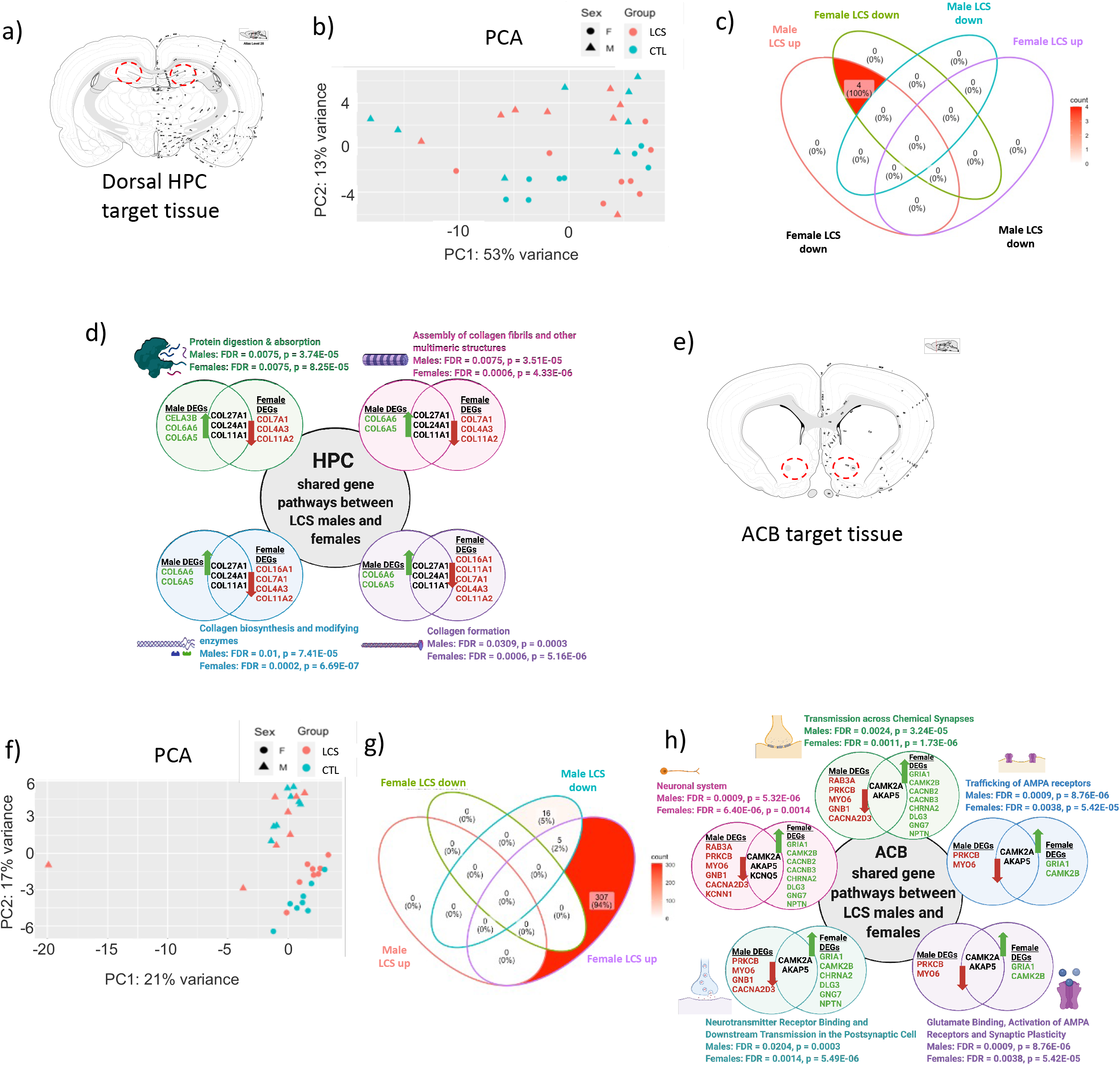
Early life LCS consumption differentially impacts gene expression patterns in the HPC and ACB. Target region for dorsal HPC tissue harvest (a). PCA of male and female LCS (acesulfame-K) and CTL rats (b). Gene pathway enrichment analyses identified 4 common gene signaling pathways that were significantly altered by LCS consumption in both males and females, as illustrated in the Venn diagram (c). All 4 of these pathways were related to collagen, and each pathway was significantly upregulated in male LCS rats, but downregulated in female LCS rats relative to controls (d). Target region for ACB shell tissue harvest (e). PCA of male and female LCS and CTL rats (f). Pathway enrichment analyses identified 5 common gene signaling pathways that were significantly altered by LCS consumption in both males and females, as illustrated in the Venn diagram (g). These pathways were related to synaptic plasticity, and each pathway was significantly downregulated in male LCS rats, but upregulated in female LCS rats. Data are means ± SEM; ns = not significant, *P < 0.05, **P < 0.01. Significant gene pathways were identified using the following parameters: p<0.05, |logFC|>=0.4 and FDR cutoff was < 0.05. CTL: control; LCS: low-calorie sweetener; HPC: hippocampus; ACB: nucleus accumbens; PCA: principal component analysis; DEG: differentially expressed gene; FDR: false discovery rate.

### 3.6 Glutamatergic plasticity-related pathways in the nucleus accumbens are altered by early life LCS consumption in a sex-specific manner

RNA sequencing analyses in ACB tissue punches (targeted region in Fig. 5e) revealed 135 DEGs for males and 227 DEGs for females overall (top 50 DEGs in ACB for each sex can be found in Supplemental Fig. 6c-d). A few of these DEGs were part of gene pathways that were altered in both males and females that consumed LCS during early life, despite the animals not having a clear separation in PCA (Fig. 5f). Gene pathway enrichment analyses revealed that several gene pathways related to glutamatergic signaling and synaptic plasticity were downregulated in LCS males but upregulated in LCS females (Fig. 5g-h). These pathways include those involved in transmission across chemical synapses (FDR = 0.0024, *P* = 3.24E-05 for males; FDR = 0.0011, *P* = 1.73E-06 for females), trafficking of AMPA receptors (FDR = 0.0009, *P* = 8.76E-06 for males; FDR = 0.0038, *P* = 5.42E-05 for females), glutamate binding, activation of AMPA receptors and synaptic plasticity (FDR = 0.0009, *P* = 8.76E-06 for males; FDR = 0.0038, *P* = 5.42E-05 for females), and neurotransmitter receptor binding and downstream transmission in the postsynaptic cell (FDR = 0.0204, *P* = 0.0003 for males; FDR = 0.0014, *P* = 5.49E-06 for females).

### 3.7 Gut microbial diversity was not affected by early life LCS consumption

We first analyzed the associations between groups (CTL/LCS) and the microbiome from phylum to ASV levels. CTL and LCS microbiomes were not separated at PCoA1 and PCoA2 for all 7 levels (Supplementary Figure 7a-g). In addition, PERMANOVA tests indicated that the microbiomes were not significantly associated with group for all 7 phylogenetic levels (*P* > 0.05). The Shannon diversity was also not significantly different between CTL and LCS groups (Supplementary Figure 7h-n). We used a linear regression model to analyze the associations between individual taxa and groups, with Group and Sex as the main effects and Group x Sex as the interaction effect. The genus *Corynebacterium.1* was the only taxa significantly more abundant in control than LCS group after adjusting for multiple testing (FDR = 0.068)) (Supplementary Figure 7o). Given that we saw effects on glucose tolerance and gene expression with early life ACE-K consumption, we analyzed the associations between the microbiome and ACE-K group separately with similar models. The PCoA ordinations, PERMANOVA tests and Shannon diversity showed similar results, with no taxonomic associations with ACE-K except that the PERMANOVA test at the phylum level indicated significant associations (*P* = 0.043) (Supplementary Figure 8). No individual taxa were significantly associated with the ACE-K group with the linear regression models after adjusting for multiple testing.

## 4. Discussion

Sugar substitution with LCS is one strategy for minimizing the detrimental effects of excess sugar intake on metabolic and neurobehavioral systems. Our findings, however, shed new light on the widespread and lasting consequences of regular LCS consumption across a sensitive developmental period spanning the juvenile and adolescence phases. Results reveal that early life LCS consumption, kept within the FDA-recommended ADI limits and consumed under voluntary conditions, significantly impairs key aspects of glucoregulation, ingestive control of caloric sugars, and memory function later in adulthood.

Even though LCS added to foods or beverages yields insignificant added calories, these compounds exert influence on nutrient intake and assimilation at various sites of action along the gut-brain axis. Here, we demonstrated that a history of daily LCS consumption during the formative stages of life leads to impaired post-oral glucose tolerance in rats, as well as specific perturbations in ingestive control for actual/caloric sugars later on during adulthood. Close inspection of consummatory patterns during a short-term intake test revealed that LCS-exposed rats were hyperresponsive to variances in the orosensory properties of two common dietary sugars, glucose and fructose, early in the ingestive episode. Furthermore, parallel analyses revealed reduced “sweet” taste receptor expression levels in the taste bud cells of LCS-exposed rats. LCS-exposed rats also displayed abnormal relative absolute levels of sugar intake in this acute intake test. That is, while rats typically consume greater amounts of glucose than fructose within a meal due to the net positive post-ingestive effects of glucose (*69*), LCS-exposed rats showed no such appetition for glucose. LCS-exposed rats were also less motivated to work for sugar in a progressive ratio task. Altogether, these data provide novel evidence that LCS consumption during critical periods of postnatal development reprograms physiological and behavioral responses to signals generated by sugars in the early phases of nutrient assimilation.

A history of habitual LCS intake early in life also affected long-term control of sugar consumption. Our results show that when LCS-exposed rats were later provided ad libitum access to a sucrose solution in their home-cage environment, they consumed more of the sugar than their LCS naïve counterparts. Our data further show that early life LCS consumption alters glutamatergic synaptic plasticity genetic pathways in the nucleus accumbens, a brain region critically involved in both appetitive and consummatory aspects of sugar reward. Further, these glutamatergic pathways were significantly affected in opposite directions by sex, with significant increases relative to controls in females and significant decreases relative to controls in males despite the fact that LCS had similar effects on behavior between sexes. Future studies will need to delineate the basis of these opposing sex-dependent effects on ACB glutamate signaling, as well as whether these motivational and ingestive perturbations stem from the heightened responsivity to the rewarding taste of sugar and/or a diminished expectation for calories conditioned by regular LCS exposure during early life.

The consequences of early life LCS consumption are not limited to ingestive control, but also include negative impacts on memory function. Previous work revealed that excessive long-term ACE-K consumption in mice led to impaired hippocampal-dependent memory function (*29*, *60*). However, these studies are limited by the fact that only one sweetener (ACE-K) and sex (males) were employed. Further, these studies were conducted in adults and involved ACE-K consumption from the only source of drinking water, which is thus involuntary and likely exceeds the FDA recommended ADI levels of daily consumption. Our study shows in adolescent animals that more limited, and voluntary exposure to LCS at the ADI level leads to hippocampal-dependent memory deficits in adulthood, as both males and females had impaired episodic memory in the NOIC task, and LCS-exposed male rats were also deficient in spatial working memory in the Barnes Maze task. Gene transcriptome pathway analyses revealed significant alterations in collagen synthesis pathways in the hippocampus following early life LCS consumption. Interestingly, despite the fact that LCS consumption was associated with HPC-dependent memory impairments in both sexes, these collagen-related gene pathway changes were sex-dependent with significant increases relative to controls in males and the opposite observed in females. Collagen plays a vital role in neural development, including in axonal guidance, synaptogenesis and glial cell differentiation (*61*), and thus the present data highlight the need for more research aimed at unveiling mechanistic links between early LCS consumption, brain collagen signaling, and neurocognitive performance.

## Acknowledgements

Funding was provided by DK123423 (Awarded to S.K. and A.F.) from the National Institute of Diabetes and Digestive and Kidney Diseases and institutional start-up funds (L.A.S.). L.T. was supported by a National Science Foundation Graduate Research Fellowship (DGE-1842487). XY was supported by DK104363.

## Contributions

L.T., L.S., and S.K. conceived the original idea and supervised the project. L.T., L.S., and S.K. designed the rat experiments. L.T., A.H., L.B., and R.L. carried out the behavioral experiments and collected samples for analyses. S.C. analyzed the taste tissue. Y.Z. and X.Y. performed RNA sequencing and gene pathway analyses on the ACB and HPCd. S.S. and A.F. analyzed the gut microbiome sequencing. L.T. prepared the figures and wrote the initial draft. L.T., L.S., and S.K. revised the manuscript with input from all authors. S.C., A.H., and E.N. contributed to interpretation, review and final editing.

---

**Supplemental Figure 1.**
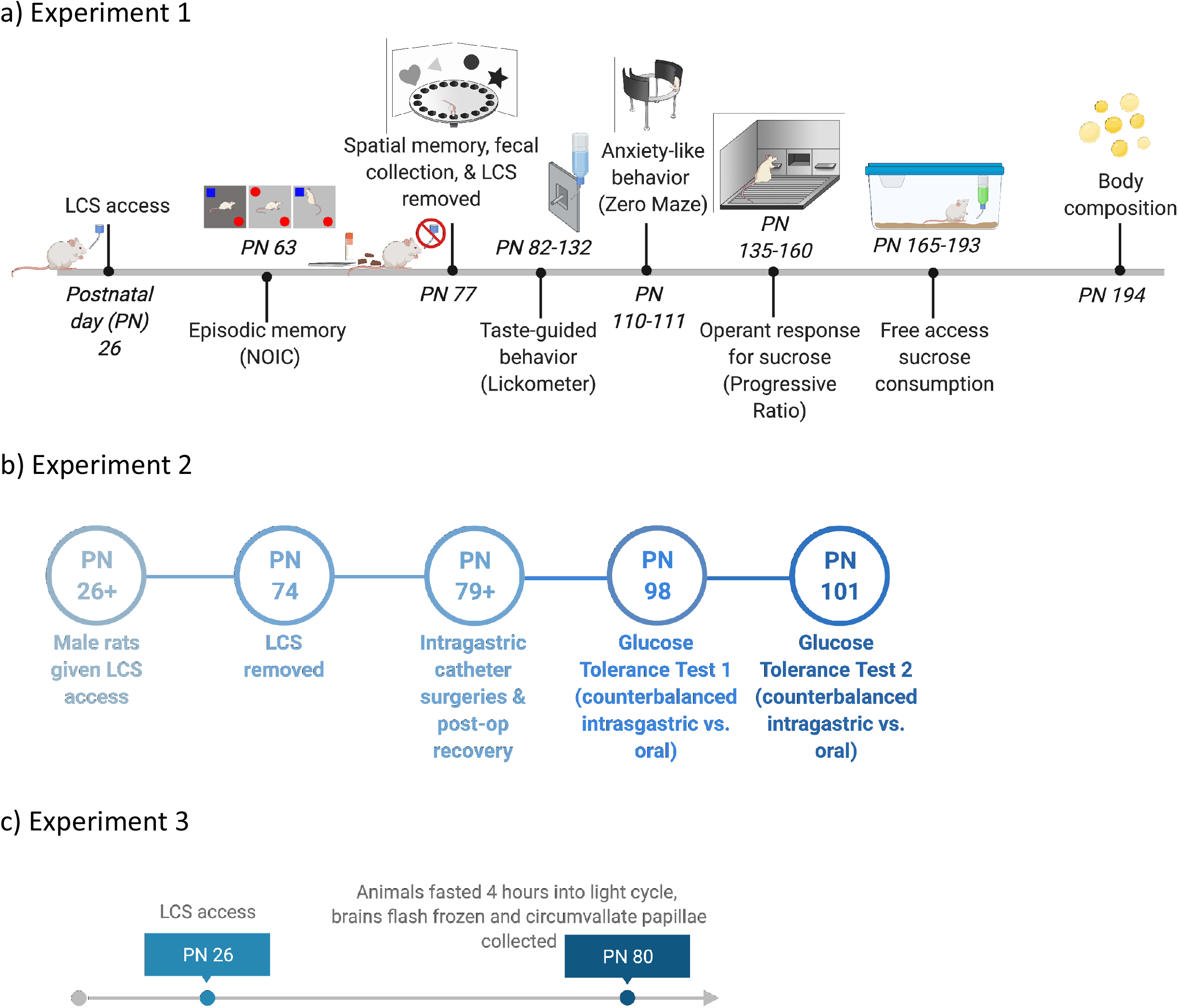
Timeline of Experiments. In experiment 1, rats were maintained on a standard rodent diet of chow and water throughout the experiment. LCS solutions (acesulfame-K, saccharin, stevia) were given daily based on mg/kg body weight doses via a sipper tube from PN 26-77. Behavioral experiments were conducted from PN 63-193 followed by body composition analysis at PN 194 (a). Timeline for the LCS access (acesulfame-K) and glucose tolerance tests in experiment 2 is shown in (b). Timeline for the LCS access (acesulfame-K) and tissue collection in Experiment 3 is shown in (c). LCS: low-calorie sweeteners; NOIC: novel object in context; PN: postnatal day

**Supplemental Figure 2.**
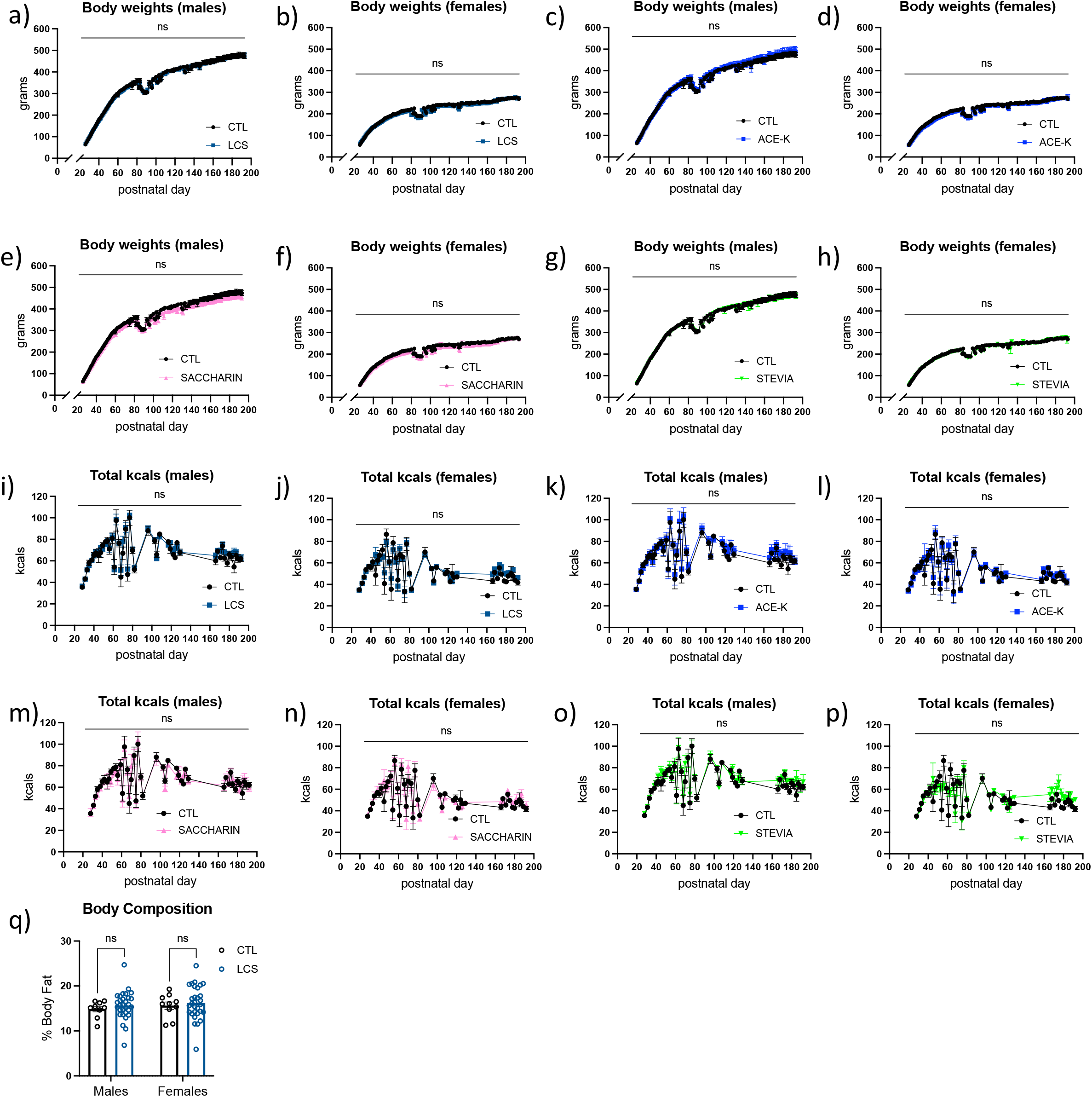
Early life LCS consumption does not impact body weight, total caloric intake, or body fat composition. Body weights for LCS combined is shown in males (a) and females (b). Body weights for males and females for each sweetener relative to CTL are shown in (c-h). Total kcals for LCS combined is shown in males (i) and females (j). Total kcals consumed for males and females for each sweetener relative to CTL are shown in (k-p). LCS consumption did not result in differences in % body fat at PN 194 (q). Data are means ± SEM; ns = not significant; ACE-K: acesulfame potassium; CTL: control; kcals: kilocalories; LCS: low-calorie sweeteners; PN: postnatal day

**Supplemental Figure 3.**
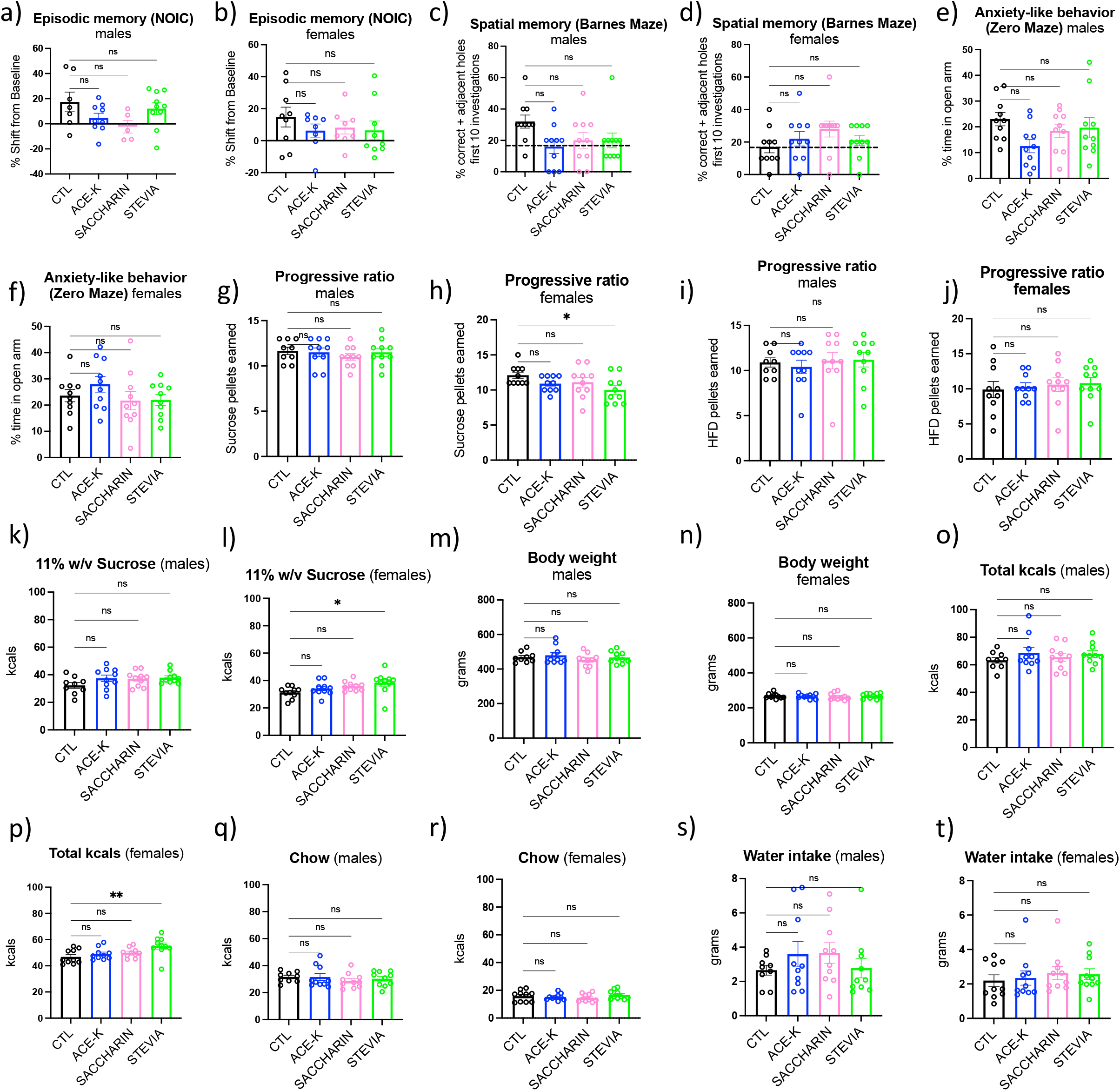
Effects of individual LCS on memory, anxiety-like behavior, and food reward-motivated behavior. No significant differences were found between individual LCS (acesulfame-K, saccharin, stevia) on episodic or spatial memory (a-d). No significant differences were found between individual LCS on anxiety-like behavior (e, f). Effects of individual LCS on pellets earned during the progressive ratio task are shown in (g-j). Effects of individual LCS on body weight and consumption during sucrose access in the home cage are shown in (k-t). Females that consumed stevia earned fewer sucrose pellets relative to CTL in the progressive ratio task (h) and consumed more calories when a 11% w/v sucrose solution was available in the home cage (p), which was driven by an increase in calories consumed from sucrose (l), but not chow (r). Data are means ± SEM; ns = not significant, *P < 0.05, **P < 0.01; ACE-K: acesulfame potassium; CTL: control; HFD: high fat diet; kcals: kilocalories; LCS: low-calorie sweeteners; NOIC: novel object in context

**Supplemental Figure 4.**
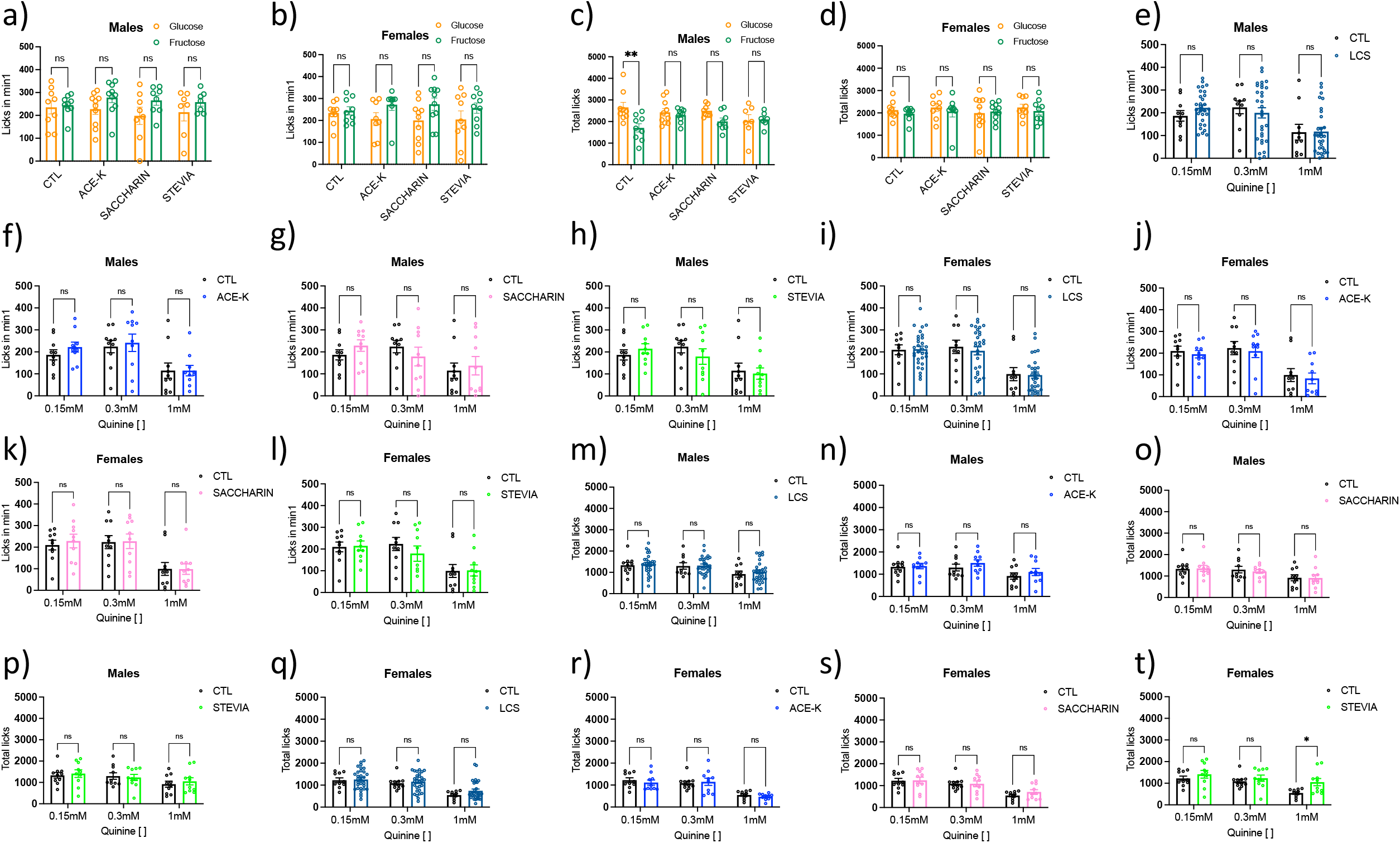
Effects of early life LCS consumption on taste-guided responding for sweet-tasting glucose and fructose and the prototypical bitterant quinine. Effects of individual LCS (acesulfame-K, saccharin, stevia) on 1^st^ minute intake and total intake over 30-minute test are shown in (a-d). Only male CTL were able to discriminate between the post-ingestive effects of glucose vs. fructose (c), whereas female CTL displayed a ceiling effect on intake (d). Effects of combined LCS and individual LCS separated by sex for increasing concentrations of quinine are shown in (e-t). Stevia females consumed more quinine than CTL at the highest concentration (1mM) over the 30-minute session (t). Data are means ± SEM; ns = not significant, *P < 0.05, **P<0.01; ACE-K: acesulfame potassium; CTL: control; LCS: low-calorie sweeteners

**Supplemental Figure 5.**
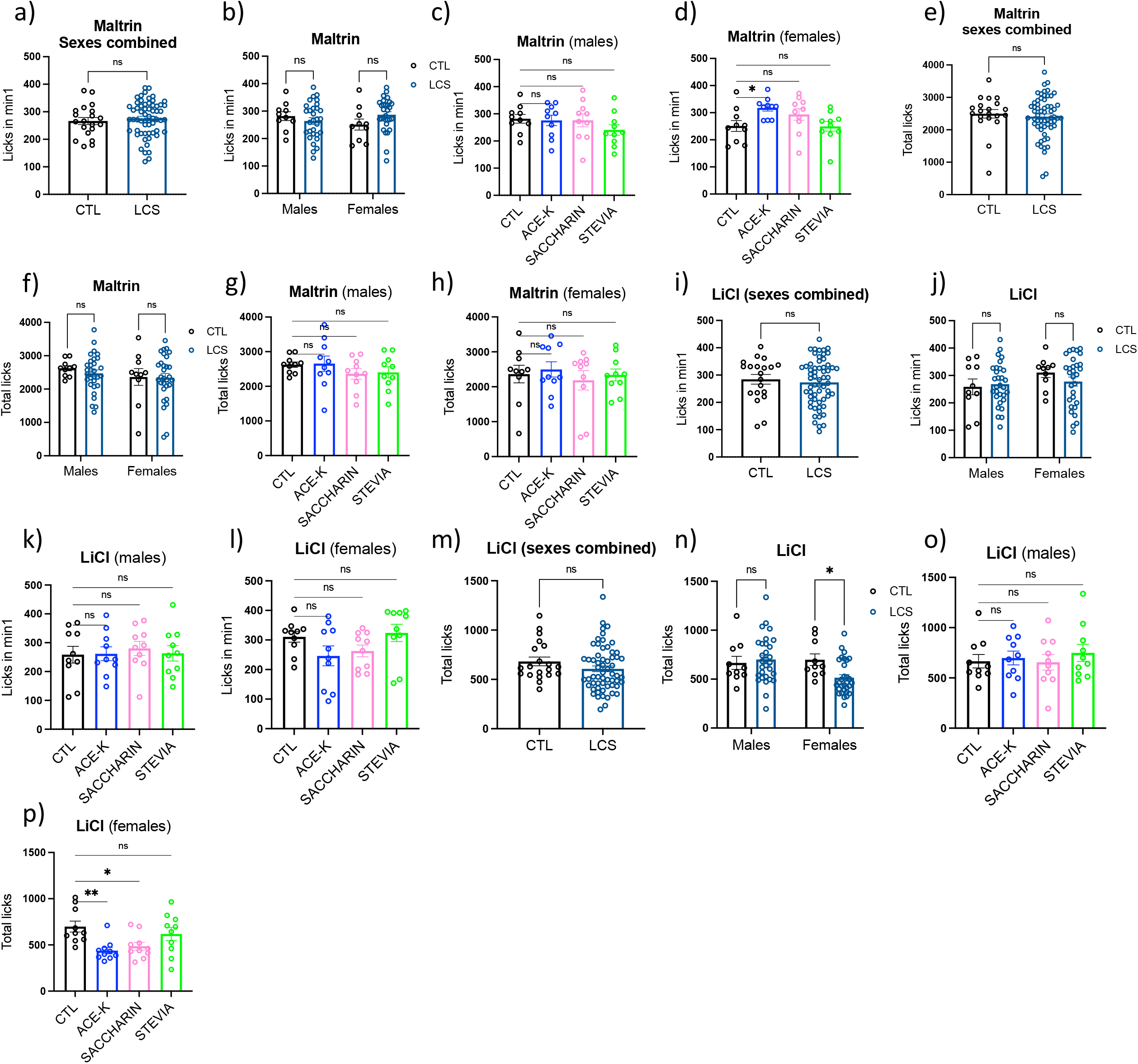
Effects of early life LCS consumption on taste-guided responding for the non-sweet carbohydrate MALTRIN and the salty lithium chloride. Effects of combined and individual LCS (acesulfame-K, saccharin, stevia) on 1st minute intake and total intake over 30 minutes for MALTRIN are shown in (a-h). Effects of combined and individual LCS on 1st minute intake and total intake over 20 minutes for lithium chloride are shown in (i-p). ACE-K females consumed more MALTRIN during the 1st minute (d), but this did not impact total intake of MALTRIN (h). LCS females were also more sensitive to LiCl, as indicated by reduced total intake in the LCS females (n), and specifically reduced intake in the ACE-K and SACCHARIN females (p). Data are means ± SEM; ns = not significant, *P < 0.05, **P < 0.01; ACE-K: acesulfame potassium; CTL: control; LCS: low-calorie sweeteners; LiCl: lithium chloride

**Supplemental Figure 6.**
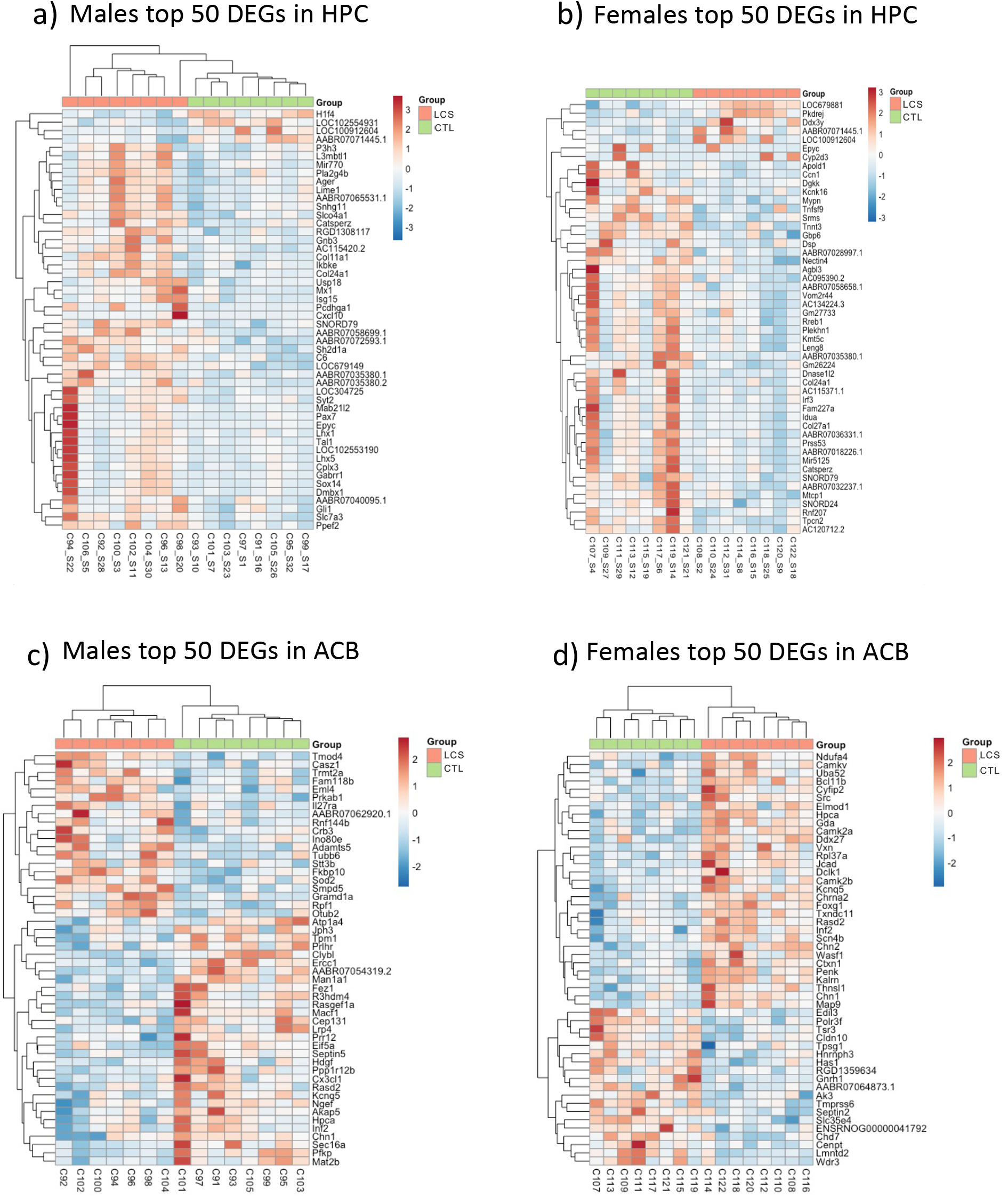
Differential gene expression analyses in the HPC and ACB following early life LCS consumption. The top 50 DEGs found in the HPC for LCS (acesulfame-K) and CTL males are shown in (a) and LCS and CTL females are shown in (b). The top 50 DEGs found in the ACB for LCS and CTL males are shown in (c) and LCS and CTL females are shown in (d). Significant DEGs were identified using the following parameters: p<0.05, |logFC|>=0.4, ACB: nucleus accumbens; CTL: control; DEG: differentially expressed gene; HPCd: dorsal hippocampus; LCS: low-calorie sweetener

**Supplementary Figure 7.**
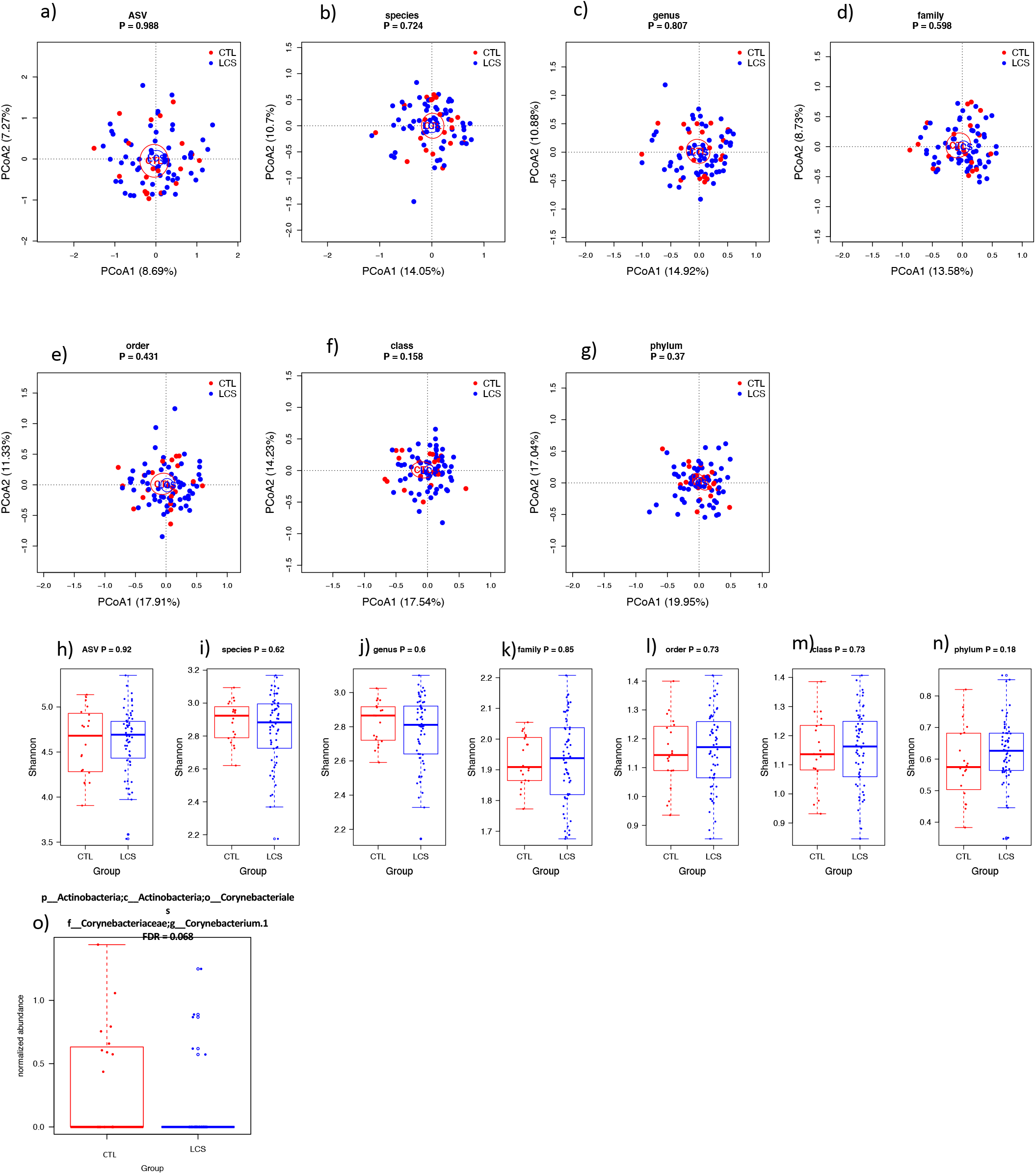
Gut microbiome analyses of CTL and LCS groups following 7 weeks of early life LCS exposure. PCoA ordinations of the microbiome of CTL and LCS groups (acesulfame-K, saccharin, stevia) at phylum to ASV levels (a-g). Ellipses indicate 95% confidence limits. P-values are from PERMANOVA tests (999 permutations). Shannon diversity of the microbiome at phylum to ASV levels (h-n). The normalized abundance (log10) of genus *Corynebacterium.1* was significantly higher in CTL (o). ASV = all species variation; CTL: control; LCS: low-calorie sweeteners; PCoA: principal coordinate analyses

**Supplementary Figure 8.**
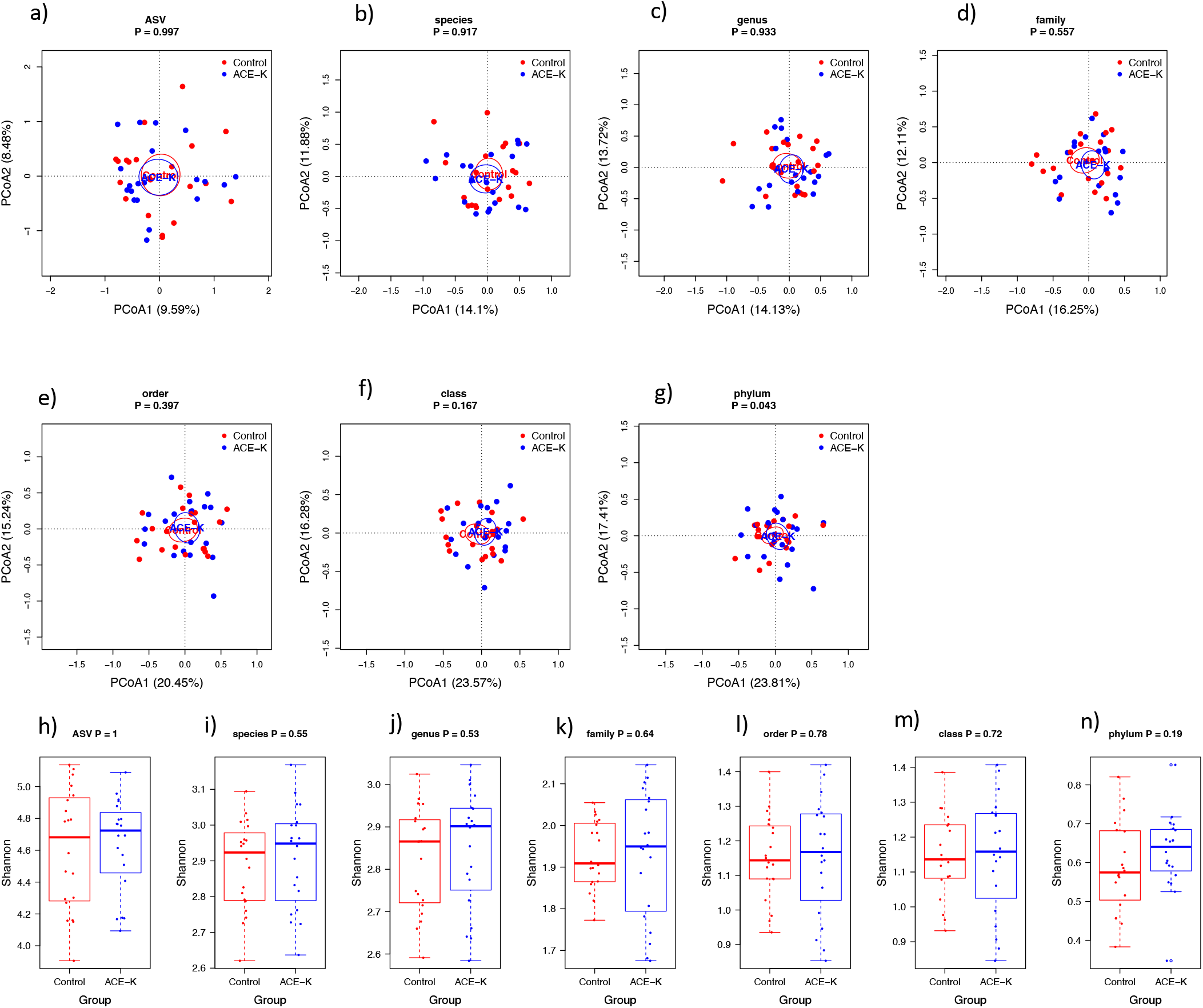
Gut microbiome analyses of CTL and ACE-K groups following 7 weeks of early life ACE-K exposure. PCoA ordinations of the microbiome of CTL and ACE-K groups at phylum to ASV levels (a-g). Ellipses indicate 95% confidence limits. P-values are from PERMANOVA tests (999 permutations). Shannon diversity of the microbiome at phylum to ASV levels (h-n). ACE-K: acesulfame potassium; ASV = all species variation; CTL: control; PCoA: principal coordinate analyses

